# Changes in gene expression in eukaryotic phytoplankton at the Atlantic-Arctic polar front

**DOI:** 10.1101/2022.11.01.514737

**Authors:** Paul Frémont, Éric Pelletier, Corinne Da Silva, Laurent Oziel, Lucia Campese, Émilie Villar, Thomas Vannier, Achal Rastogi, Jean-Marc Aury, Lee Karp-Boss, Marcel Babin, Patrick Wincker, Chris Bowler, Marion Gehlen, Daniele Iudicone, Olivier Jaillon

## Abstract

In the Arctic Ocean, phytoplankton communities, dominated by eukaryotes, sustain the marine food web and influence biogeochemical cycles. While many are endemic, some are transported by currents from the Atlantic Ocean along an environmental gradient that steepens at the polar front. Despite these distinct biogeographies, the functional characteristics of these communities remain poorly characterized. Here, we analyzed twenty metatranscriptomes from a North Atlantic–Arctic environmental gradient considering ocean currents transport times. Functions related to the regulation of gene expression and cold acclimation were more abundant in Arctic metatranscriptomes. Using the PHATE dimensionality reduction algorithm, reconstructed transcriptomes of Bacillariophyta, Mamiellales, Pelagophyceae, and Phaeocystales revealed different gene expression patterns, including photosynthesis regulation. All groups showed upregulation of numerous key temperature-associated functions and a subset of shared functions, supporting a convergence in transcriptomic changes under polar conditions. This study, based on a new resource of reconstructed transcriptomes, addressed the functional differentiation of eukaryotic phytoplankton across the North Atlantic and Arctic Ocean. It evidenced both differential and convergent changes in gene expression across the polar front in widespread phytoplankton types. These results advance our understanding of the functional dynamics of phytoplankton along environmental gradients.

## INTRODUCTION

Planktonic microorganisms thrive along global ocean currents, forming dynamic communities composed of genetically diversified populations from numerous diverse lineages of eukaryotes, prokaryotes, and viruses^1,2^. They are adapted to diverse conditions ranging from oligotrophic, warm, and saline subtropical gyres to cold, fresher, and nutrient-rich polar waters. They sustain the ocean food webs and play key roles in multiple biogeochemical processes, including primary production^3^, the biological carbon pump^4,5^ (the suite of processes that convey organic matter from the surface to the ocean interior), and nutrient recycling^6^.

Among all oceanic regions, the Arctic Ocean (AO) is one of the most extreme environments for marine life. It is characterized by extensive sea ice cover and extreme seasonality in light availability, with total darkness during the several months of polar night^7^. The AO is also a semi-enclosed basin with limited connections to the global ocean. The North Atlantic Current (NAC), representing the poleward-flowing Atlantic Waters, is the most prominent circulation feature and acts as a conveyor belt, transporting most of the heat, nutrients, and plankton toward the AO. Around 70°N, the NAC splits into two branches: one continues northward toward the Fram Strait, and the other turns eastward into the Barents Sea (Figure 1A, B). Together, the Fram Strait and Barents Sea represent the main gateways of the AO, accounting for 80% of total inflow and outflow. The NAC therefore plays a pivotal role in structuring ecological connectivity between the NAO (North Atlantic Ocean) and the AO.

**Figure 1.**
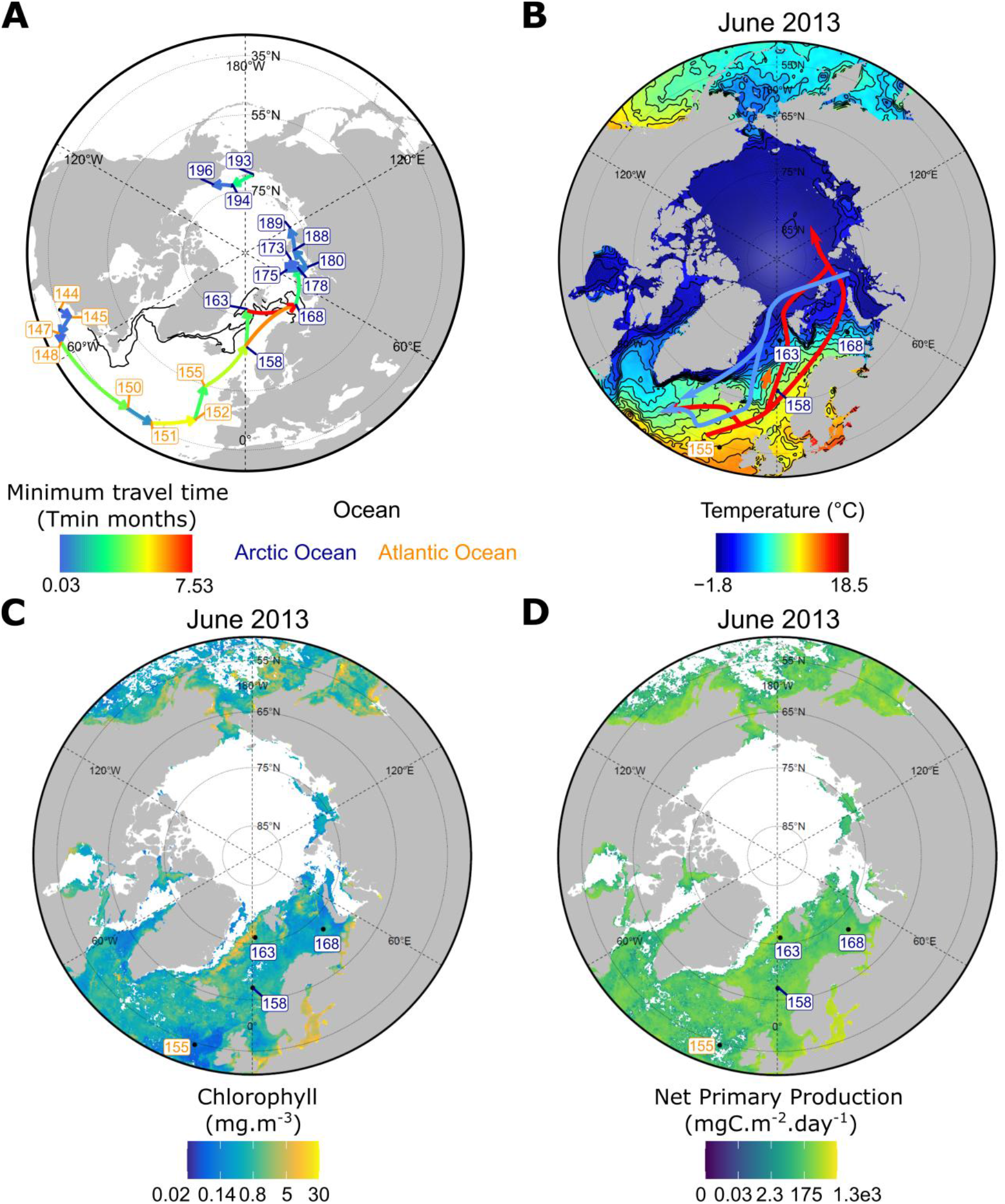
Physical connectivity by currents of studied samples and environmental context of the study. (**A**) *Tara* Oceans and *Tara* Oceans Polar Circle stations considered in this study, and the shortest Lagrangian path (minimum travel time Tmin) between stations, starting at Station 144. (**B**) Mean Sea Surface Temperature in June 2013 around the time of sampling of stations 155 to 168 (between May and July). The orange arrow indicates the polar front between Stations 158 and 163 where a steep temperature gradient was observed (black lines). Main inflow and outflow currents are represented by red arrows (warm NAO inflows) and blue arrows (cold Arctic outflows). (**C**) Mean Chlorophyll concentration in June 2013 around Stations 155 to 168. (**D**) Mean Net Primary Productivity (NPP) in June 2013 around Stations 155 to 168. Chlorophyll and Net primary productivity were computed using an ocean color algorithm developed for the AO^41,42^ (Methods). NPP is interpolated from chlorophyll concentrations, resulting in some missing values being filled, while other areas are not interpolated due to masking by sea ice.

While being transported into the AO from the NAO mostly *via* the NAC, the community structure and diversity of many plankton groups have been shown to drastically change^8–15^. Many physico-chemical parameters change along this route although temperature has been shown to be the primary driver of community structure. Some studies have suggested that the transition between temperate and polar communities would occur at latitudes around a thermal threshold in the range of 10-15°C^8,10^. A principal community feature of AO phytoplankton is its predominance of eukaryotic groups over cyanobacteria^16–21^, a trend that contrasts with that observed in other marine basins. This pattern is principally attributed to the higher maximal growth rate of larger cells which allow eukaryotic phytoplankton to outcompete smaller cells in high nutrient conditions^22^. Using 18S taxonomic annotations, sampling stations from the *Tara* Oceans Polar Circle expedition have already confirmed a strong differentiation between AO and NAO eukaryotic communities^13^. However, several major eukaryotic phytoplankton lineages appear to be present in both basins. Among these are small protists, picoeukaryotes such as Mamiellales, nanoeukaryotes like Phaeocystales, and taxa of larger organisms like diatoms and dinoflagellates^13,16–18,21^.

The AO is currently experiencing one of the fastest rates of warming of the planet^23^ warming at a speed up to four times the global average^24^. The European sector of the AO is also subject to an increasing influence of Atlantic waters from the NAC, reduced sea ice cover, increased ventilation and disrupted stratification, in a process often referred to as “Atlantification”^25^. Simultaneously, these processes are modifying plankton community structure, with the appearance of temperate dinoflagellates and coccolithophores^26,27^, and the dominance of small sized picophytoplankton in certain parts of the AO at the expense of large-celled phytoplankton such as diatoms (Bacillariophyta)^28^. These changes are particularly affecting plankton phenology and ecology, raising questions about the resilience of Arctic ecosystems^23^.

The environmental change that plankton communities are experiencing during their transport to the AO motivates the need to understand the related physiological response of plankton communities. The context of the amplified climate change in the AO region further stresses this question. The physiological response to environmental changes in plankton has started to be experimentally studied for some individual taxa^29– 31^, and some studies have begun to address metabolic differentiation using environmental omics data^8,10,32–39^. According to previous reports, bacteria^8^ and eukaryotes^10^ have overall different biological activity between the AO and temperate regions. Notably, polar eukaryotes have an enrichment in functions linked to catalytic activity acting on proteins and RNAs^10^ and translation is correlated with temperature^39^. For individual taxa, several studies focus on diatoms^19,36,37^. For example, in the southern ocean, local adaptation, both in gene expression and at the sequence level has been shown in the diatom *Fragilariopsis cylindrus* comparing light and dark conditions, temperature and low iron^19^. In diatoms, cold-shock proteins and the unknown domain DUF285 with a structure similar to anti-freeze proteins, are more transcribed in polar regions^36^. Specific diatoms also change their physiology in bloom conditions in the Antarctic with increased energy production and response to oxidative stress^37^. In the Pelagophyceae clade, one species changes its gene expression involving stress oxidative responses, lipid metabolism and photosystems in response to sudden salinity decrease^38^.

Here, we perform a comparative metatranscriptomic analysis of eukaryotic-enriched plankton samples collected at 20 locations situated in the NAO and AO, to test three hypotheses. First, the strong physical connectivity driven by ocean currents should generate a detectable biological continuum across the basins. In addition, the polar front is expected to act as an environmental filter, imposing selective pressures that would shape community composition. Secondly, we hypothesize that the interplay of these two forces lead to progressive genomic and functional differentiation, with the strongest effects occurring at the polar front.

Third, we hypothesize that different lineages might show differential gene expression patterns due to phylogenetic divergences but also convergent responses along the same environmental gradient.

To test these hypotheses, we analyzed deeply sequenced metagenomic and metatranscriptomic datasets, aligned at 95% nucleotide identity, using a comprehensive unigene catalog^36^. For the first hypothesis, we compared minimum transport times by currents with metagenomic and metatranscriptomic data. For the second hypothesis, we used the taxonomic and functional annotation of the gene catalog focusing on more abundantly transcribed functions in the AO. For the third hypothesis, based on co-abundance patterns of unigenes across sampling stations, we assembled a collection of metagenomics-based transcriptomes (MGTs), each of which would encompass multiple unigenes from a specific lineage. To assess the physiological changes of these MGTs, we used a gene expression metric that normalizes metatranscriptomic abundances by metagenomic abundances.

## RESULTS

### Oceanographic context

As part of *Tara* Oceans, the *Tara* schooner conducted plankton sampling at eight sites in the NAO from January to March 2012, followed by sampling at twelve sites in the AO from May to September 2013 (*Tara* Oceans sampling sites 143 to 196, as depicted in Figure 1A). This approach ensured a seamless progression across the sampled seasons. Calculated from prior investigations^40^, the minimum transport time by currents (Tmin) between adjacent stations (adjacency being defined by this minimal transport time) revealed a prevailing current trajectory from the NAO into the AO (Figure 1A, Figure S1 and Figure S2). En route to the AO, the vessel initially took the western route via the Greenland Sea, passing south of the Fram Strait (*Tara* Oceans stations 158 to 163, Tmin=3.2 months). Later, the vessel diverged from the main current, transitioning to the Barents Sea route (*Tara* Oceans stations 163 to 168, Tmin=7.5 months; stations 158 to 168 indicated by the orange arrow, Tmin=6.1 months). Beyond station 168, diverse AO regions were sampled, primarily influenced by NAO inflows, except for stations 193 to 196 which were notably affected by Pacific inflows (Figure 1A). The polar front is discerned through a pronounced temperature gradient observed during the sampling period (June 2013) between stations 158 and 163 (orange arrow in Figure 1B). We further characterized each sampled station using their Temperature/Salinity (TS) at sampling and other different depths, distinguishing the two sets of stations (AO and NAO) and polar front stations (Figure S3 A, B). In addition, 3 to 4 water masses could be assigned to the different sampling points (Figure S3C). Finally, there was an increase in chlorophyll concentration (Figure 1C) and net primary production (Figure 1D) as the vessel traversed into the AO region.

Given this marked shift in environmental conditions and the associated change in planktonic primary production, as well as other likely biological processes along the NAC, we initially aimed to characterize the genomic differentiation of eukaryotic communities across the two basins.

### Genomic differentiation between the Arctic and North Atlantic Oceans

In a previous analysis, this sample collection has already been described from a taxonomic perspective using 18S ribotypes^13^. From this analysis, only 10 to 16% of eukaryotic plankton ribotypes were shown to be strictly endemic to the AO and in low abundance while 9 to 12 % were found to be polar indicators and were much more abundant (Figure 2 and 3 from Ibarbaltz *et al*.^13^). Half of the relative abundance of eukaryote ribotypes corresponds to photosynthetic groups, among which 80% were comprised of the following four groups: Bacillariophyta (10 to 80%), dinoflagellates (Dinophyceae) (10 to 45%), Haptophyceae (2 to 10%), and Mamiellophyceae (<1 to 2%). Therefore, a significant taxonomic differentiation exists between the NAO and AO communities.

**Figure 2.**
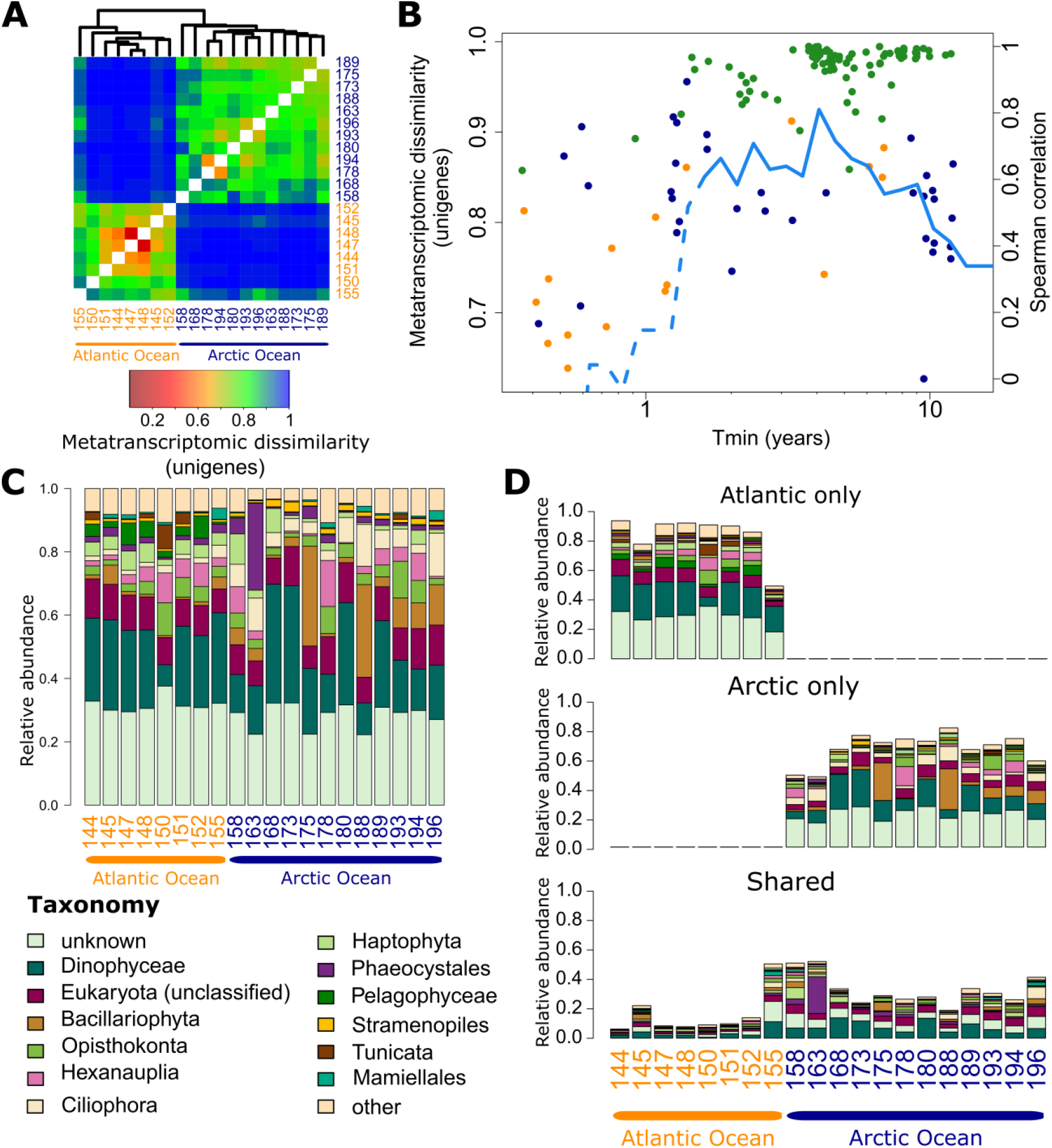
Genomic differentiation of the Arctic Ocean (AO) and North Atlantic Ocean (NAO), its relation to oceanic connectivity and composition of metatranscriptomes. (**A**) Hierarchical clustering of the matrix of metatranscriptomic pairwise β diversity estimates based on unigene relative abundances (Bray-Curtis dissimilarity index, Methods). (**B**) Metatranscriptomic diversity as a function of Lagrangian connectivity between pairs of *Tara* Oceans and *Tara* Oceans Polar Circle stations (minimum transport time by currents) and cumulative Spearman correlation coefficient between the two metrics (blue line). When the line is full, the correlation is significant (p<0.05). Orange, dark blue and green points respectively indicate NAO-NAO, AO-AO, and NAO-AO pairs of stations. (**C**) Metatranscriptomic taxonomic composition based on relative abundances of unigenes from major plankton taxa (taxonomic level can vary) in the NAO and AO (stations ordered by increasing sampling number). (**D**) Decomposition of the taxonomic composition between basin-specific unigenes and shared unigenes between the two basins (classified based on presence or absence of expression). A clear transition enriched in shared unigenes appears in Stations 155, 158 and 163.

**Figure 3.**
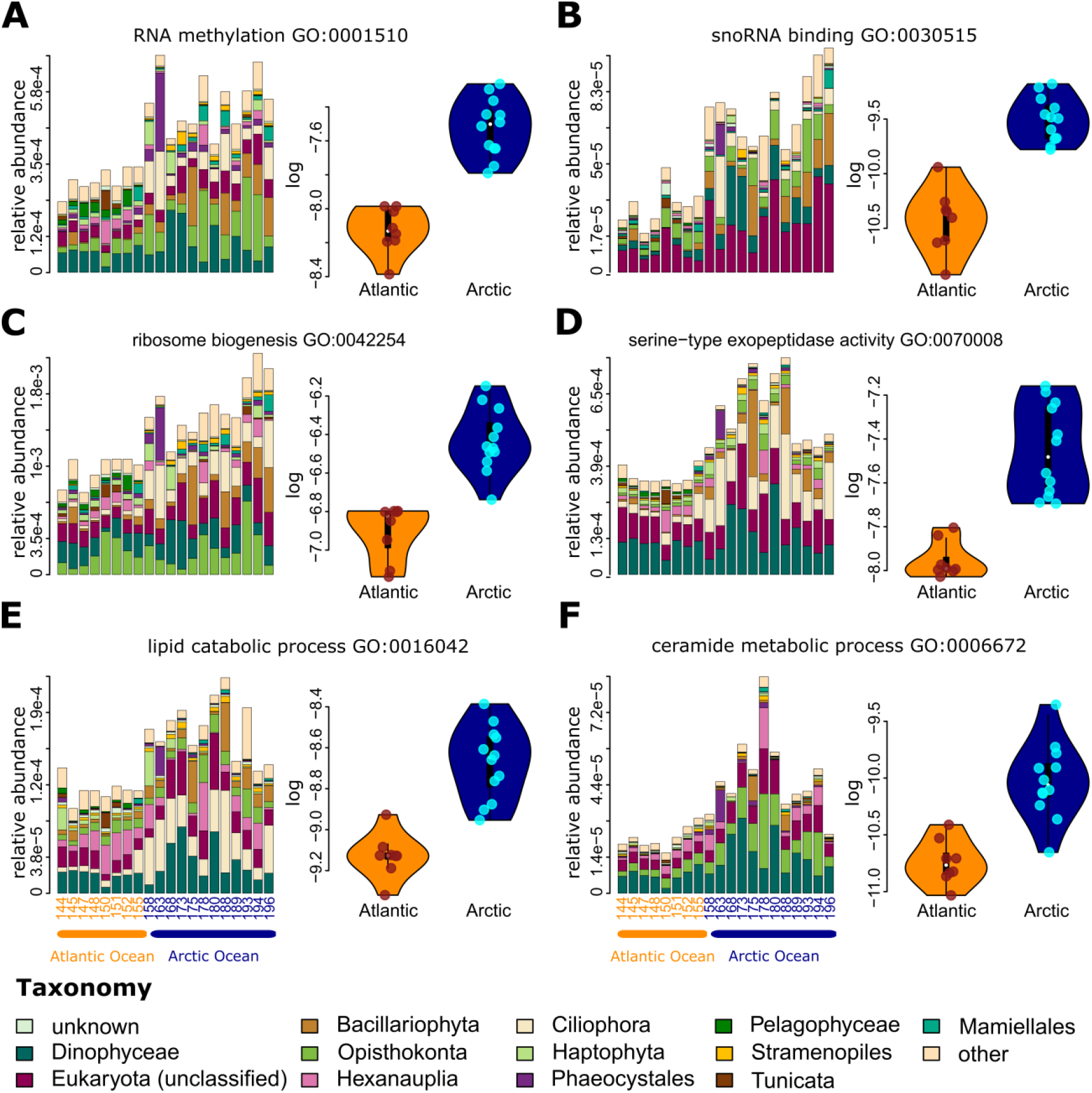
Examples of more abundant key functions in Arctic Ocean (AO) metatranscriptomes. Each panel (**A**-**F**) represents (left) the sum of the metatranscriptomic relative abundances of unigenes associated with the specified Gene Ontology (GO) terms across the considered taxa and (right) the distribution of this relative abundance in the NAO (orange) versus AO (dark blue). Each of these GO terms is significantly more abundant in the AO (t-test, Hommel correction p<0.05). The comprehensive list is available in Table S1.

Here, we used a raw dataset comprising 0.6 Tb of metatranscriptomic and 0.6 Tb of metagenomic sequences to investigate the biogeographical structure (Figure S1). We analyzed changes in plankton community composition through two forms of β diversity (βD) across metagenomes and metatranscriptomes of the 0.8-2000 µm size fraction (Figure 2, Methods, equations 4 and 5). The first approach, relying on DNA fragments, has been demonstrated to corroborate ribotype-based results and has been previously analyzed at an ocean-scale^40^. The second utilized a recently published unigene catalog^36^. Considering the samples for this study, βD is computed from the relative abundances of approximately 69 million unigenes (Figure S1) within this catalog. These two estimates of βD exhibited strong correlations (Methods). The βD assessments revealed both a substantial differentiation between the AO and NAO and an inherent internal heterogeneity within each basin (Figure 2A). This heterogeneity in the AO did not align with distinct oceanographic regions^13^. Based on the βD matrix clustering (Figure 2A), we categorized stations 143-155 as NAO and stations 158-196 as AO (Figure 1A and 2A). Developing a genomic “atlanticity” index for each station based on the βD (Methods), the atlanticity index aligned with the TS pattern continuum of the stations at sampling depths (Figure S3D). The temperatures of the first transition stations (155-163) towards the AO were located between 10 and 15°C which is close to previously reported break point in eukaryotic βD^10^.

Based on our genomic definition of the AO and NAO, 10% (n=6.9 million) of expressed unigenes from the catalog were common to both basins, whereas 55% (n=38.2 million) were unique to the NAO and 35% (n=24.1 million) were unique to the AO (Figure 2 C, D). Based on the relative abundances of the different taxa (Figure 2, S4), significant differences were found for Ciliophora and Pelagophyceae and other less abundant groups between the two basins (Figure S5). The relative abundances of common unigenes revealed a clear transition between the two basins corresponding to the polar front (Figure 2D). Stations 155, 158 (NAO side of the front) and 163 (AO side of the front) were found enriched with common unigenes (51.2% of the read abundance and 42.3 % of the unigenes in number on average, Figure 2D, Figure S4 and S6 for unigenes counts). This outcome suggests the mingling of diverse communities originating from distinct water masses, a well-documented phenomenon in oceanic fronts^20^.

### Link between physical connectivity and genomic differentiation

To assess the role of surface ocean circulation on βD, we compared the minimum transport times (Tmin) separating sample sites with the corresponding βD estimates (Figure 2B, Figure S7A). A maximal correlation (r=0.75, p<0.05) emerged around 4 years of Tmin and remains higher than 0.6 between 1.5 and 8 years of Tmin, aligning with the temporal intervals typically separating NAO and AO stations (light blue dots in Figure 2B). A similar slightly lower correlation was also found between an environmental distance (ED) and Tmin (Figure S2C). The correlation between βD and Tmin suggest that a current-driven influence on community composition, previously measured in the NAO with a maximum correlation at 1.5 years^40^, extends poleward. We further characterized the intricate interplay between the physico-chemical environments, current and plankton communities by visualizing environmental differentiation of the samples (Figure S2A), minimum transport times (Figure S2B) and the relation between βD and the environmental distance during transport (Figure S7B, C).

### Metatranscriptome-scale functional changes in the polar environment

The clear environmental differences between the AO and NAO, coupled with substantial variations in the composition of their eukaryotic plankton communities, suggest divergent functioning. To investigate this question, we leveraged the Gene Ontology (GO) functional annotations of the unigene dataset. We computed the metatranscriptomic relative abundance of expressed unigenes carrying these GO terms (see Methods) to establish an operational metric that reflects the relative contribution of a specific function within community transcripts. This metric integrates both organism abundances and transcription levels. Our statistical analysis identified from 66 to 529 out of 4,695 terms significantly more abundant in the AO metatranscriptomes (illustrative examples in Figure 3) and 11 to 230 in the NAO (see Methods). The comprehensive list of functions displaying significantly higher abundance in both basins and both metatranscriptomes and metagenomes is available in Table S1 (t-test, Hommel correction p<0.05, see example of more abundant GOs in NAO metagenomes in Figure S8).

From the reduced set of 66 GO terms more abundant in AO metatranscriptomes, 20 were related to RNA processing, while 11 were associated with RNA methylation, including 6 to tRNA methylation (Figure 3A, Table S1). The importance of RNA methylation in cold stressed plants has been studied previously^43^. As an example, one of the significantly more abundant terms, ‘tRNA (adenine-N1-)-methyltransferase activity’ is known to be important for stabilizing tRNA^44^ and participating in mRNA methylation and transcriptome protection within cytoplasmic granules under stress^45^. ‘tRNA methylation’ is also recognized for its role in translational fidelity^46^. Finally, adverse conditions can alter the balance between free tRNA and aa-tRNA^43^, crucial for accuracy of protein synthesis^47^.

Other significantly enriched GO terms in the AO were tied to gene expression regulation and downstream processes. These included terms related to regulation of gene expression exemplified by ‘snoRNA binding’ (Figure 3B), ‘ncRNA processing’, ‘mitochondrial RNA metabolic process’. Notably, ncRNAs, including snoRNAs and mitochondrial RNA metabolism are pivotal for gene expression regulation^48,49^. snoRNAs engage in RNA chemical modification, including methylation, while ncRNAs (small RNAs) have a vital role in cold acclimation in *Arabidopsis*^50^. Additionally, terms like ‘ribosome biogenesis’ (Figure 3C), ‘preribosome, small subunit precursor’, and ‘ribonucleoprotein complex biogenesis’ were significantly more abundant. Ribosomes serve as cold and heat shock sensors^51^, with their dynamics and stability altered under cold conditions^52^. Functions like ‘protein peptidyl-prolyl isomerization’, ‘peptidyl-proline modification’, ‘serine-type exopeptidase activity’ and ‘serine-type carboxypeptidase activity’ (Figure 3D) were also more abundant in the AO. *Lato sensu*, these functions could be associated with post-translational modifications.

We also identified other functions that were more abundant in the AO, such as chromosome integrity maintenance and various functions linked to lipid catabolism (lipid catabolic process’ (Figure 3E) and ‘fatty-acyl-CoA binding’), membrane fluidity (‘Ceramide metabolic process’ and ‘Sphingolipid metabolic process’), protein transport and organelle membranes organization (Table S1). Functions related to protein trafficking (protein transmembrane transport’ and ‘mitochondrial inner membrane presequence translocase complex’) and enzymes activities (‘serine-type exopeptidase activity’ and ‘serine-type carboxypeptidase activity’) were more abundant in the AO. In contrast, functions linked to cellular compartments protein retention, such as ‘maintenance of location’, maintenance of protein localization in organelle’ and ‘protein retention in ER lumen’ were found more prevalent in NAO metagenomes (Figure S8F, Table S1). These results hint at an accelerated protein turnover in spring/summer AO plankton communities.

We additionally used the ALDEx2 package^53,54^ to compare PFAM abundances between the AO and NAO. Interestingly, almost all PFAMs related to cold stress and acclimation (Table S2) were significantly more abundant in the AO in both metagenomes and metatranscriptomes (Figure S9). Particularly, the ice binding domain (PF11999) stands out as the most excessively abundant both at the metatranscriptomic level and within specific taxa (approximately 215 times higher in the AO).

Although not ruling out seasonal effects and acknowledging some variations within the AO (Figure 3 D, B, E, F for instance), these patterns appeared throughout the entire spatiotemporal sampling of the AO basin (from May to September, as shown in distribution plots of Figure 3).

In summary, comparing AO and NAO metatranscriptomic data confirmed significant transcriptional changes in the AO for eukaryotic plankton. Notably, these changes involved gene expression regulation tied to cold acclimation and gene expression regulation.

### Changes in gene expression patterns

In the previous sections we have analyzed metatranscriptomic abundances, which are strongly influenced by species abundances and community composition. To better capture relative gene expression changes across samples, we normalized transformed metatranscriptomic abundances by corresponding metagenomic data for each PFAM and each unigene at each station (Methods and Figure S10).

First, based on PFAM expression levels, we defined a gene expression distance (methods) following previous work in prokaryotes on the same plankton samples to compare among stations the rates of change in community composition with that of gene expression^8^. Like what was reported for prokaryotes, we observed a very clear modification below 15°C with a greater contribution of community turnover with respect to gene expression differences (Figure S11).

Second, we aimed to characterize, at the level of individual genes, patterns of gene expression regulation which could reflect a response to the environmental change. To this end, we analyzed the gene expression patterns of approximately 572,000 unigenes that were present in both metatranscriptomes and metagenomes from at least 5 stations. We used the PHATE^55^ (Potential of Heat diffusion for Affinity-based Transition Embedding) dimension reduction technique to create a 2-dimensional embedding of the expression matrix for these 572,000 unigenes (Figure 4) (see Methods). PHATE aims to capture global and local structure as well as probabilistic transitions in the data space and visualize them in a reduced number of dimensions^55^. We used the Euclidean distance as the metric for the PHATE embedding to account for gene expression values.

**Figure 4.**
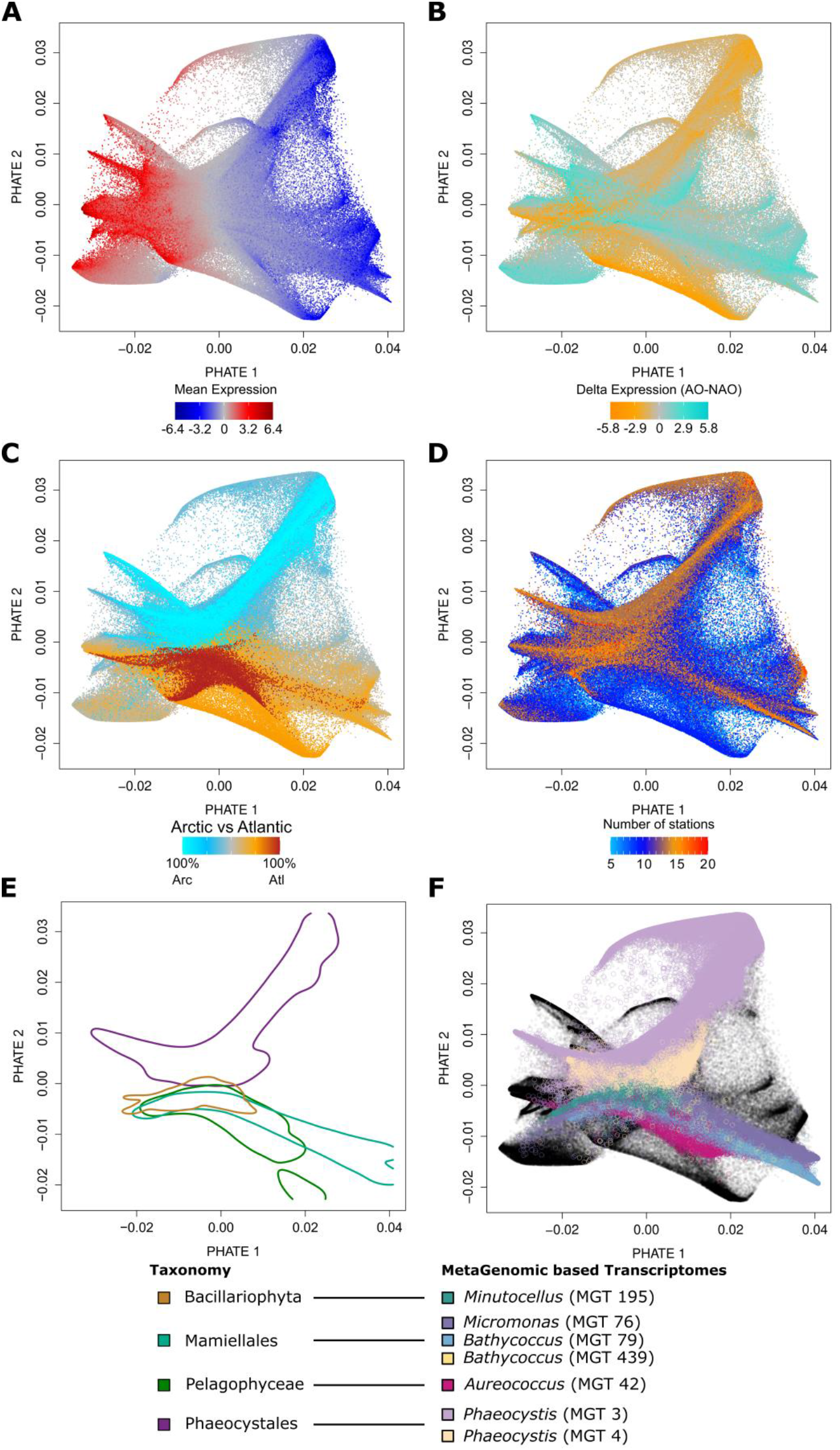
Topological patterns of the two dimensional embedding of the gene expression matrix. Each plot shows the same PHATE embedding of the expression matrix of approximately 572,000 unigenes expressed in at least 5 stations. Each point represents the position of the expression profile across the 20 samples of one unigene. Each unigene is colored according to different biological and ecological characteristics: (**A**) Mean level of expression across samples (CLR). (**B**) Mean delta of expression between the AO and NAO. (**C**) An index characterizing its biogeography (Methods). (**D**) The number of stations where unigene expression was detected. (**E**) Unigene density contours for the main eukaryotic phytoplankton groups analyzed. (**F**) Seven widespread eukaryotic metagenomics-based transcriptomes (MGTs) are colored in the PHATE space.

The result of this 2-dimensional embedding visually presented a distinct yet intricate structure (Figure 4). Initially, we explored whether specific environmental and/or biological parameters available to us could account for this observed structuring. The first axis (PHATE 1) displayed a strong correlation with the mean expression values of unigenes (r=0.89, p-value<0.001, Fig. S12A), evident in the embedding visualization (Figure 4A). The embedding was also patterned by the mean expression differences between the two basins (Figure 4B), albeit with a less clear association to the PHATE axes. This suggests that the expression of many genes was regulated along the transition between the NAO and AO. The second axis (PHATE 2) exhibited a notable correlation with the biogeography of the unigenes, representing their degree of endemicity or cosmopolitanism across the two basins (r=-0.57, p-value<0.001), as discernible in the embedding (Figure 4C). The number of stations where unigenes were expressed and the standard deviation of their expression were also identified as secondary drivers of the structure (Figure S12). Looking at the contribution of each station, the biogeographical structure was also visible with NAO stations mainly in the bottom part, transition stations covering wider space and AO stations mainly the top part (Figure S13). Moreover, some stations explained that some of the branches of the embedding likely correspond to functional and taxonomic idiosyncratic features (that can be shared by several stations) (Figure S13).

To further analyze the PHATE space, we used a previously described procedure^56^ to group unigenes based on their co-varying abundance across metagenomic data. This approach is analogous to the construction of metagenome-assembled genomes^57^ (MAGs) but here, the genes are assembled using metatranscriptomics in a first step and secondly binned using metagenomes (see Methods for details). This procedure generated approximately one thousand metagenomics-based transcriptomes (MGTs), likely representing gene clusters from the same species/strain. Among this collection, seven phytoplanktonic MGTs were widespread and abundant within the considered size fraction and the 20 samples analyzed. These MGTs were taxonomically associated to major groups of eukaryotic phytoplankton, including two *Bathycoccus* (Mamiellales), one *Micromonas* (Mamiellales), two *Phaeocystis*, one *Aureococcus* (Pelagophyceae), and one *Minutocellus* (a pennate diatom). Together they represented ∼136,000 unigenes (24% of the 572,000 unigenes expressed in at least 5 stations). In this study, we manually curated the taxonomic annotation of these MGTs by using the name of the closest available reference. Looking at the distribution of each phytoplankton group within the PHATE space, they occupied varied and generally distinct areas, aligning with different behaviors in the communities (Figure 4E). For the regions associated to Mamiellales and Phaeocystales, MGTs occupied a specific subspace (Figure 4F) further clarifying patterns in the PHATE space. The presence of taxonomically closely related MGTs situated in proximity likely reflect transcriptomic and niches divergences. Particularly Mamiellales, Pelagophyceae and to a lesser extent Bacillariophyta unigenes were more predominant in the NAO while Phaeocystales were more predominant in the AO (Wilcoxon test p<0.01).

The six MGTs showed overall average gene expression levels which were significantly different from each other (Figure 5A, pairwise Wilcoxon test p<0.01, from the most to the least expressed: Bacillariophyta MGT 195, Phaeocystales MGT 4, Pelagophyceae MGT 42, Phaeocystales MGT 3, and Mamiellales MGT 76, 79 and 439). All considered MGTs except that from Bacillariophyta had an overall gene expression lower than the community average (Wilcoxon test, p<0.001, Figure 5A). MGTs 195, 76, 79, 439 and 4 were found to be significantly more active in the AO than in the NAO while MGT 3 was more active in the NAO (Wilcoxon test p<0.001, Figure 5C). Zooming on photosynthesis genes, including photosystems, Light Harvest Complexes and Cyclic Electron Transport (PFAMs considered in Table S3), photosynthesis was generally a significantly highly expressed metabolism and was significantly more expressed in the Phaeocystales MGTs than in the Bacillariophyta and other MGTs (Figure 5B). Finally, photosynthesis was generally less upregulated than average in Mamiellales in the AO while generally more upregulated in the AO in Phaeocystales (Figure 5D). Visualizing the raw data, photosynthesis expression seemed more heterogeneous than average gene expression and was very highly expressed in some stations with a less clear link to the basin (Figure S14A vs B). In particular, the upregulation of photosynthesis in the AO for Mamiellales and Bacillariophyta MGTs comes from one or two stations and a few more stations for the Phaeocystales MGTs.

**Figure 5.**
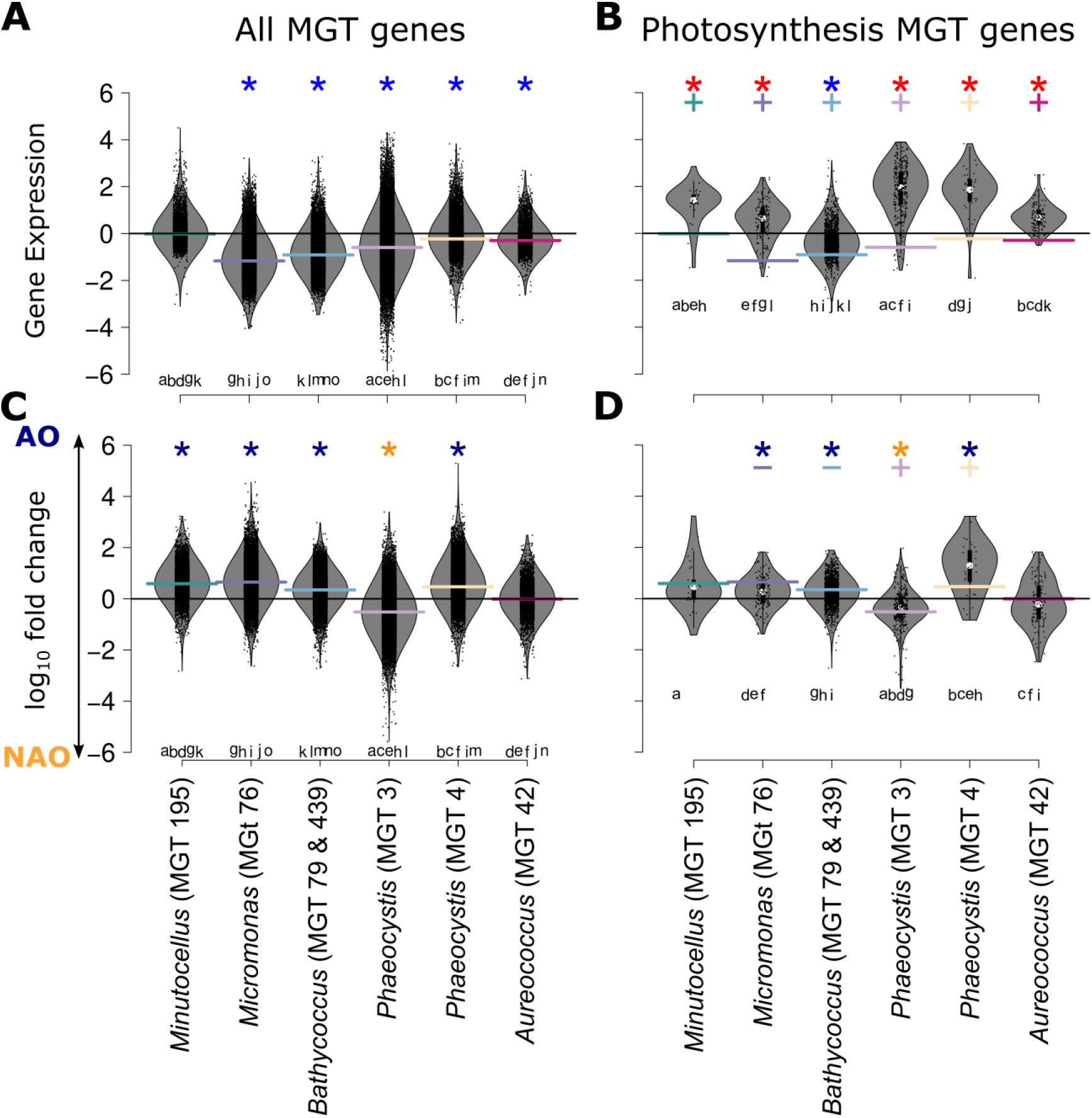
Distribution of unigene expression levels from six metagenomics-based transcriptomes (MGTs) and their differential expression between the Actic Ocean (AO) and North Atlantic Ocean (NAO). Violin plots shows the mean gene expression levels for six MGTs from major phytoplankton groups, for (**A**) all unigenes and (**B**) photosynthetic unigenes. Blue and red stars indicate medians significantly lower, or larger than the mean, respectively (µ!=0, Wilcoxon test, p<0.001). Panels (**C**) and (**D**) show the difference in mean expression levels between AO and NAO for the same MGTs and gene sets. Dark blue and orange stars mark medians significantly lower or higher than zero (µ!=0, Wilcoxon test, p<0.001). Colored plus and minus signs indicate medians significantly larger or lower, than the mean delta of each MGT (represented by colored horizontal bars). Letters at the bottom of the plots indicates statistically significant pairwise comparison (pairwise Wilcoxon test, p<0.01).

Overall, these results suggest differential activities as well as regulation of photosynthetic activity among phytoplankton groups and basins.

### Convergent gene expression shifts at the polar front

We postulated that within the abundant groups of phytoplankton in both basins the expression of certain gene functions might be altered in a similar manner along the environmental gradient. To explore this hypothesis, we used a subset of approximately 485,000 unigenes, present and expressed in both basins, from the previous 572,000 that we analyzed (Figure S1). We examined their expression values in relation to the physical environmental parameters and further scrutinized their functional annotations.

We identified approximately 88,500 instances of significant correlations, positive or negative, between gene expression profiles of unigenes and specific physical parameters among temperature, sunlight duration (SSD), and salinity. While these physical environmental parameters were strongly intercorrelated, temperature emerged as the parameter significantly correlated with the expression profiles of a larger number of unigenes (Figure S15, ∼19,500 negative and ∼12,700 positive, FDR=0.05).

By investigating the unigenes whose expression appeared negatively correlated with temperature (indicating an increase in expression from NAO to AO) we found that a particularly large number of them were from diverse groups of picophytoplankton, particularly Mamiellales, Bacillariophyta and Pelagophyceae (Figure 6A, Figure S15). Therefore, we performed a more detailed analysis of the taxonomic and functional distribution of these unigenes. Across all tested phytoplankton taxa, unigenes with expression levels negatively correlated with temperature exhibited strong overlap in their functions (GO terms) and appeared clustered in a reduced dimension space (apart from all positively correlated gene repertoires, Figure 5B-C). These results are summarized by comparing the proportion of unigenes sharing GO terms among the different groups, separately for positive and negative correlations with temperature. On average this fraction was significantly higher for unigenes negatively correlated with temperature among the phytoplankton groups (Figure 5D).

**Figure 6.**
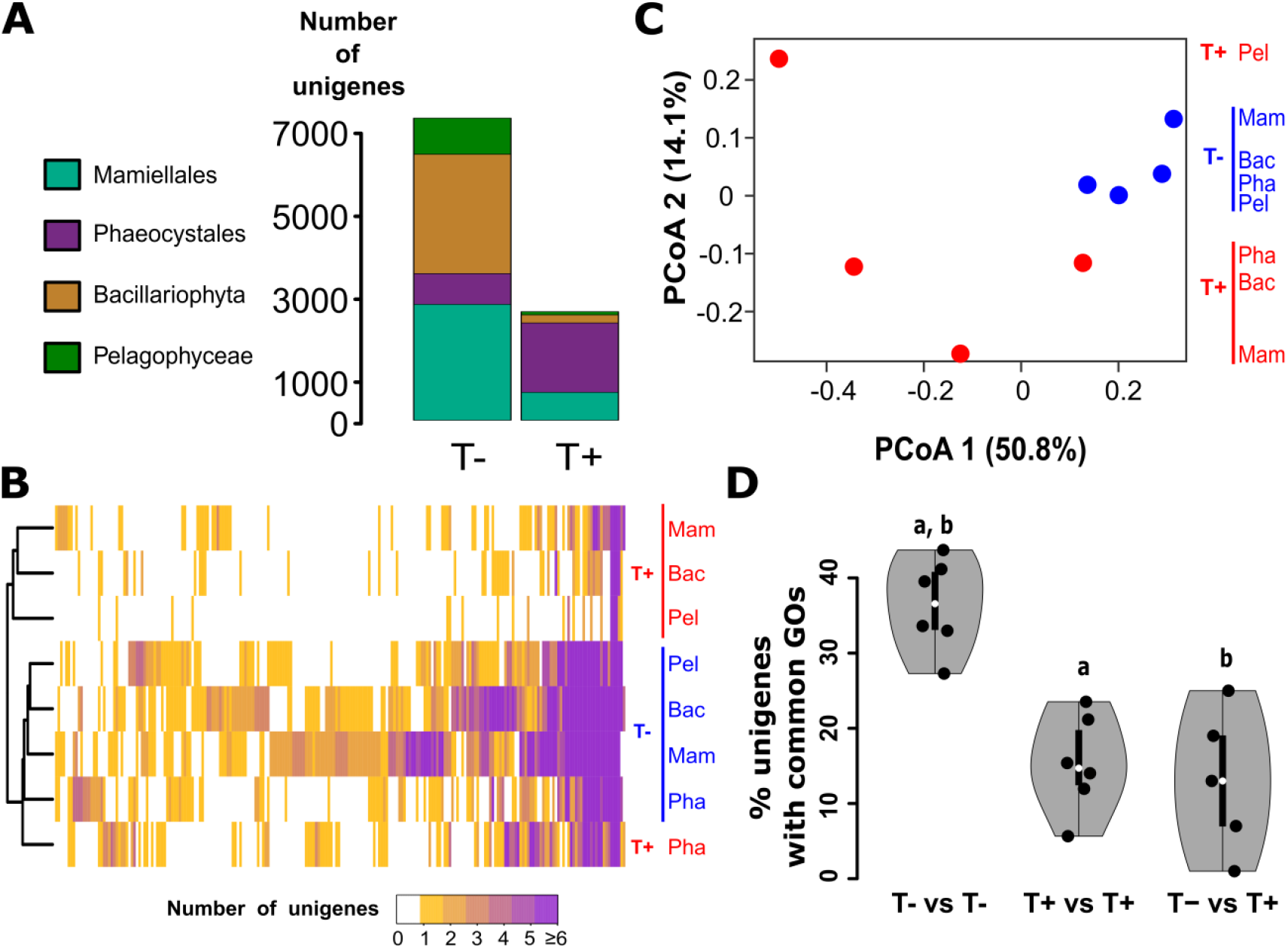
Convergent gene expression changes in eukaryote phytoplankton that traverse the polar front. (**A**) Taxonomic decomposition of unigenes whose expression profiles are significantly correlated with temperature, either negatively (left bar) or positively (right bar). (**B**) Heatmap of the number of unigenes and associated Gene Ontology (GO) terms that correlate with temperature across the main taxa. Each row represents the number of unigenes associated with a GO term (in column) for a given taxa and a correlation sign (blue: negative, red: positive). Rows are clustered based on the Bray-Curtis dissimilarity index. (**C**) Bray-Curtis-based Principal Coordinate Analysis (PCoA) of panel (**B**). (**D**) Distribution of the percentage of unigenes with common GO terms for unigenes either positively or negatively correlated with temperature among phytoplankton groups. Each point represents a comparison for a pair of phytoplankton groups. Letters indicate significant pairwise differences (Wilcoxon test, p<0.01, holm correction).

These findings suggest that the temperature cline encountered by phytoplankton at the NAO to AO transition leads to a transcriptomic response in a set of core metabolic functions within these groups. This regulation appeared similarly across various lineages regardless of their phylogenetic relationships. Beyond this core set of functions, other functions were also seen to exhibit the same response only in certain lineages (Figure 6B). We found many such functions in Mamiellales, Bacillariophyta and Pelagophyceae, but fewer in Phaeocystales. We noted that Mamiellales and Bacillariophyta had the most similar cold-responsive gene repertoires, reminiscent of the functional convergence in their gene repertoires recently revealed in MAGs^58^.

Within the MGTs, we found significant enrichments of functions previously described to be linked to cold acclimation, light response, or osmotic stress (Figure 7, Figure S16 for other MGTs and Figure S17 for group level analysis). In MGT 79, *Bathycoccus* (Figure 7A), a unigene involved in the thiamine diphosphate (Vitamin B1) biosynthetic process was negatively correlated with temperature. This process is known to be important for phytoplankton nutrition and growth^**59**^ but also in early-stage response to cold and osmotic stress in Arabidopsis^**60**^ and poplar^**61**^. In MGT 439, *Bathycoccus* (Figure 7A), the trehalose biosynthetic process was an enriched GO term in negatively correlated unigenes linked to cold acclimation^**62**^. In MGT 195, annotated as the pennate diatom *Minutocellus* (Figure 7B), the expression profiles of a unigene involved in tocopherol cyclase activity (also in MGT 42 *Aureococcus*, Figure S16C) and another in the allantoin catabolic process were significantly negatively correlated with temperature. Tocopherol cyclase activity is important for cold^**63**^, salt^**64**^, and high light acclimation^**65**^ in plants. Allantoin, a metabolic intermediate of purine catabolism, often accumulates in stressed plants^**66**^ and enhances cold stress tolerance in rice grains^**67**^. In MGT 4, a *Phaeocystis* (Figure S16C), superoxide dismutase (SOD) (also in MGT 439, *Bathycoccus*, Figure 7A), heat shock protein binding, and glutamyl-tRNA reductase activity are enriched GO terms in negatively correlated unigenes. SOD increases stress tolerance in potato through mitochondrial activity^**68**^ and is also important in wheat^**69**^ and barley^**70**^. Heat shock proteins respond to cold stress in *Synechococcus*^**71**^ and some plants^**72**^. Glutamyl-tRNA reductase, a key enzyme for 5-aminolevulinic acid (ALA) synthesis, is highly expressed in cold stress response and remains at high levels after acclimation, although this varies across plants^**73**^. Phospholipase A2, which plays a role in cold acclimation in wheat^**74**^, is enriched in MGT 76, *Aureococcus* (Figure S16A). At higher taxonomy levels (Figure S17), other functions associated with cold stress were also enriched, such as tocopherol cyclase activity^**63–65**^, pantothenate biosynthesis^**75**^ in *Pelagophyceae*, and polyketide metabolic process^**76**,**77**^ in *Dinophyceae*. All these functions whose expression are negatively correlated with temperature and previously described to be important in cold acclimation in terrestrial plants and aquatic organisms, suggest shared strategies of gene expression changes across the green lineage. In addition, the enriched functions included processes related to numerous functions related to gene expression regulation, energy consumption, DNA division/replication, motility, nucleotide biosynthesis, protein folding, DNA repair and unknown functions (Figure 7 and S16-17).

**Figure 7.**
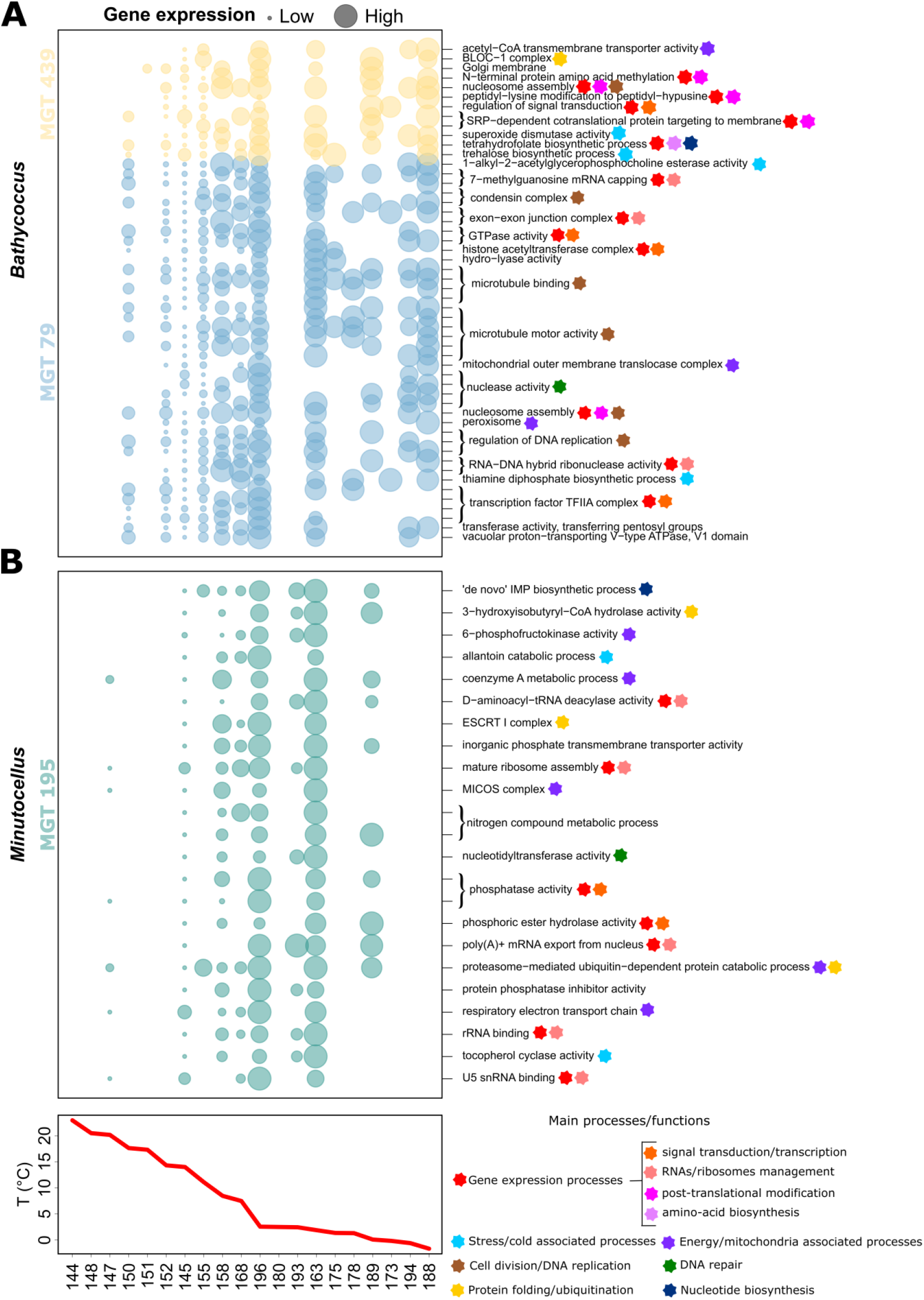
Expression profiles and functional annotation of enriched unigenes negatively correlated with temperature for two Mamiellales and one Bacillariophyta metagenomics-based transcriptomes (MGTs). Expression profiles (left) of unigenes whose Gene Ontology terms (right) are enriched within unigenes significantly negatively correlated with temperature are displayed for two phytoplankton taxa from three metagenomics-based transcriptomes (MGTs): (**A**) *Bathycoccus* (MGT 79 and 439 with the same taxonomic annotation) and (**B**) a pennate diatom (MGT 195, annotated as *Minutocellus*). For this MGT, unassigned functions (unknown) were enriched; however, they were too numerous to be represented here. Empty spaces indicate that no expression data is available. Expression values (represented by the bubble sizes) are scaled so that the maximum expression value is equal for each unigene. Stations are ordered according to the temperature gradient shown at the bottom.

## DISCUSSION

In this study, by leveraging a recently published collection of unigenes and a novel collection of metagenomics-based transcriptomes (MGTs) we have characterized structural and functional changes in a substantial portion of the eukaryotic component of planktonic communities occurring along a major current zone that transports them from the NAO into the AO. These analyses were conducted at three levels related to our initial hypotheses: the impact of current connectivity on biogeographic structuring, functional differences between the two basins, and gene expression changes in major eukaryotic phytoplankton groups.

As described in previous reports^8,10,11^ our data confirm a strong genomic and transcriptomic differentiation between polar and North Atlantic communities. Previous studies used diverse sequencing approaches (metabarcoding, metagenomics, metatranscriptomic) either from the samples we used here^8,11^ or from other expeditions^10^. In agreement with these studies^8,10^, we found a transition from temperate to polar plankton communities around 10-15°C.

In the line of our previous work on temperate regions^40^, our data support the hypothesis that strong physical connectivity would generate a biological continuum across the basins, despite the environmental filtering effect of the polar front. We obtained a significant correlation between β diversity and the estimated minimum transport time by currents. This correlation extended between 1.5 and 8 years of transport time, longer than the previously reported 1.5 year continuum observed in other basins^40^. This underscores the lasting influence of ocean currents on the composition of communities at genomic resolution, even when confronted with abrupt environmental changes around the polar front. Importantly, this influence is observed for our samples that cover different years and seasons which suggest the maintenance of global structure as previously proposed^40^. Additionally, numerous MGTs and unigenes were present in both basins, suggesting that similar or connected populations were maintained across the transition zone. Given that we used a 95% nucleotide identity threshold, which is commonly used to delineate genomic units^56,78,79^, although debated for the definition of species^80^, these may correspond to different populations most likely from the same species, or potentially from very closely related species. Whether these populations physically cross the front or are independently maintained on both sides remains unresolved. A population-genomics-based report using the same nucleotide threshold on metagenomics sequences mapping the Mamiellales *Bathycoccus prasinos* suggested a biogeographical structuring clearly distinguishing polar populations^81^. Other studies suggest that certain diatoms can regulate buoyancy^82^, offering a plausible mechanism for connection between populations and persistence across fronts, and water masses.

Following our second hypothesis and extending the results from Martin *et al*.^10^, we further characterized significant functional disparities between the sampled communities distinguishing the two basins. Notably, many functions related to gene expression regulation *lato sensu* (from signal transduction to post-translational modification) were found to be more abundant in AO metatranscriptomes. These functions included processes related to RNA methylation, proposed to be important for maintaining RNA stability^44^, ensuring precise protein translation^46,47^ and contributing to cold stress responses^43^. These findings indicate that many post-transcriptional mechanisms may be activated in this polar environment. In contrast to Martin *et al*.^10^ that associated ribonucleoprotein complexes with warm water PFAM networks, we found that ribosomal biogenesis and ribonucleoprotein complex synthesis were generally more abundant in AO metatranscriptomes. This apparent discrepancy may stem from differences in the regions or seasons sampled, as our NAO dataset was collected in winter. In addition, in our sample these functions were more abundant only in metatranscriptomes and not in metagenomes, supporting a regulatory origin.

Our third hypothesis proposed that lineages found in both basins would primarily exhibit distinct gene expression profiles but paradoxically also exhibit convergences for some functions related to the environmental gradient. We observed lineage-specific transcriptional differences for Mamiellales, Bacillariophyta, Pelagophyceae and Phaeocystales, among others. Using a two-dimensional embedding of unigene expression, we detected structured patterns based on expression levels, basin identity, and taxonomy. These patterns appeared for large taxonomic levels but were further refined down to the resolution of MGTs. Although the cell size of organisms was not considered here, this might be an important parameter in gene expression levels^83,84^. Interestingly, while *Phaeocystis* MGTs were the most prevalent taxa in the AO in our dataset, the most expressed MGT was annotated as the diatom *Minutocellus* which also have the largest cells (among considered taxa). Diatoms are believed to be the most important primary producers, important for carbon cycling in the Arctic^85^ and dominant in blooms^86^. Yet, *Phaeocystis* MGTs show evidence for a higher photosynthetic activity in this taxon and throughout a larger number of stations in the AO than the Diatom MGT (and others). Interestingly, an increasing photosynthetic activity of *Phaeocystis* has been reported in the Antarctic via a symbiosis by cytoklepty of an Acantharian species^87^. Although we did not find evidence for this symbiosis in our data, the high photosynthetic activity of these Phaeocystis MGTs was compatible with these reports and their key role in carbon cycling^88^.

Our third hypothesis proposed also that distantly related lineages would share similar changes of expression level for some functions along the same environmental gradient. Temperature was the environmental parameter most correlated with transcriptomic variation of a wide panel of unigenes across multiple lineages. A subset of eukaryotic phytoplankton found in our data exhibited a similar transcriptomic response with a substantial number of genes from Mamiellales, Bacillariophyta, and Pelagophyceae. Evidently, this correlation does not imply a causal relationship between temperature and gene expression. Nevertheless, these gene expression changes involved various cold acclimation-related functions^59–77^. They included many functions found in other organisms of the green lineage, highlighting their conservation. Other metabolisms negatively correlated with temperature included gene expression regulation functions (from signal transduction to post-translational modification), energy consumption processes, protein folding/ubiquitination, cell division/DNA replication, nucleotide biosynthesis, and DNA repair. Notably, as discussed above, considering only the transcriptomic abundance (and not expression levels), functions linked to gene expression regulation were also present at higher levels throughout the whole AO samples (covering the end of spring and the whole summer).

It is important to acknowledge several methodological limitations, most inherent to the meta-omics approach. First, we focused exclusively on the 0.8–2000 μm size fraction, which is enriched in pico- and nano-phytoplankton. This likely underrepresented phytoplankton groups having large cells, such as some diatoms which can have a non-negligible role in phytoplankton biomass through local blooms^86^. Second, despite a relatively high shotgun sequencing effort (30 GB for each of the twenty metatranscriptomes and each of the twenty metagenomes), our gene catalog was probably biased toward the most abundant species^89^. Yet, following a previously described methodology^56^, this effort allowed us to reconstruct a new collection of one thousand MGTs likely representing widespread species and with compact genomes. Therefore, Dinoflagellates, known for their large genomes^90^, were missing from the MGTs. Finally, while changes in transcript abundances were often used as proxies for metabolic activity, the relationship between gene expression and actual physiological rates remained indirect. Although previous studies reported strong correlations between gene expression and environmental conditions, yet in other contexts^32,33^, this link may not apply uniformly for all metabolic functions and taxa. Despite these limitations, the depth of our dataset, the taxonomic breadth of the MGTs reconstructed, and the clear functional signals observed across the NAO-AO environmental gradient provided robust support for the three hypotheses tested in this study.

Our results raise important questions about the mechanisms underlying gene expression plasticity. Plankton populations, generally with large sizes, are composed of numerous genotypes^91^. Each genotype has an acclimation range, which is the ability at the individual level to change gene expression appropriately in response to environmental change. Accordingly, genotype-level adaptation, which involves changes in protein sequences that are beneficial for the given environment and vertically transmitted to next generations, has been observed in certain cases, such as *Bathycoccus*^81^ and the SAR supergroup^92^, as well as through mechanisms like allele-specific expression reported in the copepod *Oithona similis*^93^ and the diatom *Fragilariopsis cylindrus*^19^. The patterns we report here are most likely from populations of cells that, in addition to showing different gene expression patterns, may also carry genomic polymorphisms potentially linked to adaptation to the Arctic environment. It will be of interest to disentangle these two aspects in future studies. Investigating both transcriptional and sequence level changes at a broader scale and for more organisms will be critical to better understand what factors differentiate the widespread and relatively generalist lineages that we have studied here, from more local and specialist ones^94^, their response to a changing climate, as well as consequences on ecosystems^23,95^.

In the context of rapid climate change, these two mechanisms —acclimation and adaptation—are expected to modulate future community structures. Adaptation might require more time^96^ but can occur relatively fast in plankton with short generation times and large population sizes^94,97^. Moreover, the strong connectivity between the AO and NAO by the North Atlantic Current^98^ is expected to intensify these processes in the context of global warming^26^ and as a consequence of continued “Atlantification” of the Arctic^25^.

In conclusion, this study has improved our understanding of eukaryotic phytoplankton gene expression and functional changes across the NAO/AO environmental gradient. Our results indicate both lineage-specific and convergent gene expression changes to the environmental transition. In the context of Atlantification of the AO it is important to assess which species have the potential to adapt and acclimate to future conditions.

## LEAD CONTACT

Further information and requests for resources should be directed to and will be fulfilled by the lead contact, Paul Frémont.

## ETHICS APPROVAL AND CONSENT TO PARTICIPATE

Not applicable.

## CONSENT FOR PUBLICATION

Not applicable.

## FUNDING

Paul Frémont was supported by a CFR doctoral fellowship and the NEOGEN impulsion grant from the Direction de la Recherche Fondamentale of the CEA. This study received funding from the European Union’s Horizon 2020 Blue Growth research and innovation programme under grant agreement number 862923 (project AtlantECO).

## AVAILABILITY OF DATA AND MATERIALS

All code and data used to reproduce the figures are available at https://github.com/PaulFremont3/metaT_AO_NAO (pre-release v1). Large.rds files necessary to reproduce the figures, including the results of gene expression correlations with environmental parameters, the ALDEx analysis, unigenes taxonomy annotation, and NOAA Sea Surface Temperature, can be downloaded from https://doi.org/10.5281/zenodo.17316416. Contextual data are available in Pangaea (DOI: 10.1594/PANGAEA.875582) and a simplified version is included in the github repository. The NOAA OI SST V2 High Resolution Dataset is available at https://psl.noaa.gov/data/gridded/data.noaa.oisst.v2.highres.html. The Remotely Sensed Chlorophyll a and Net Primary Production Data is available at http://albedo.stanford.edu/gertvd/research/arctic/prod/. Files for the Marine Atlas of Tara Oceans Unigenes version 1.5 (MATOU-v1.5) and for the metagenomics-based transcriptomes (MGT-v1.5), including FASTA sequences, taxonomic and functional annotation tables, and abundance tables, are available at https://www.genoscope.cns.fr/tara/. Metagenomic and metatranscriptomic raw data are deposited at the European Nucleotide Archive under accession numbers PRJEB402, PRJEB9691, PRJEB9738, and PRJEB9739.

## ACKNOWLEDGMENTS

We thank the LAGE (Laboratoire d’Analyses Génomiques des Eucaryotes, CEA) members for stimulating discussions on this project, the commitment of the Research Federation for the Study of Global Ocean Systems Ecology and Evolution (FR2022/TaraGOSEE) and of Stazione Zoologica Anton Dohrn, Alessandro Tagliabue for providing iron PISCESv2 model data, Joshua S. Weitz for his commentaries on the manuscript, C. Scarpelli and members of the scientific computation team from Genoscope for support on computations. We thank all members of the *Tara* Oceans consortium for maintaining a creative environment and for their constructive criticism. *Tara* Oceans would not exist without the *Tara* Ocean Foundation and the continuous support of 23 institutes (https://oceans.taraexpeditions.org/). This study benefited from access to high-performance computing resources through GENCI-[TGCC/CINES/IDRIS] and the ESPRI computing and data centre (https://mesocentre.ipsl.fr), which is supported by CNRS, Sorbonne Université, Ecole Polytechnique and CNES and through national and international grants. This article is contribution number XX of *Tara* Oceans.

## AUTHOR CONTRIBUTIONS

Conceptualization: OJ, DI, MG, LK, CB, PW, PF, TV; methodology: PF, OJ, EP, CD, JMA, EV, LO, LC, AR, TV; investigation: PF; writing—original draft: PF, OJ; writing—review & editing: PF, OJ, MG, DI, CB, EP, LO, MB, PW; funding acquisition, OJ, PW, DI, MG, EP; resources, EP, JMA, CDS; supervision: OJ;

## DECLARATION OF INTERESTS

Authors declare no competing interests.

## DECLARATION OF GENERATIVE AI AND AI-ASSISTED TECHNOLOGIES

During the preparation of this manuscript, the authors used ChatGPT to assist with grammatical corrections and language clarity. After using this tool, the authors reviewed and edited the content as needed and accept full responsibility for the content of the publication.

## METHODS

### MATOU v1.5

We used a recently updated version of the Marine Atlas of *Tara* Oceans Unigenes that included metatranscriptomic samples encompassing the AO^36^. This extension includes a total of 581 samples (140 more than the previous version which did not include Arctic samples) from 6 different size fractions divided between 0.8 and 2000 μm. Each sample was sequenced to an average depth of 30 Gbp following the methodology of Carradec *et al*.^32^. In total, all samples were collected at 88 locations across the oceans (previously 68) at three depths (Surface (SRF), Deep Chlorophyll Maximum (DCM) and Mesopelagic zones (MES)). We followed the same methodology as previously described for the sequencing, assembly, clustering, mapping, taxonomic and functional annotation for the updated atlas. This catalog totals 158.3 million non-redundant transcribed sequences, called unigenes, increasing by 41.7 million the previous version (+35.7%). In this study we used a subset of 63 million unigenes from the catalog from size fraction 0.8-2000 µm and encompassing the 20 AO and NAO surface samples (Figure S1). This dataset embraces many eukaryotic phyla covering both phytoplanktonic and non-phytoplanktonic groups (Figure 2).

### Metagenomics-based transcriptomes (MGT v1.5)

We built a novel version of the metagenomics-based transcriptomes (MGTs), based on the approach originally described by Vorobev *et al*.^56^. MGTs group unigenes based on their relative abundances’ covariations across metagenomic samples. The MGTs are clusters of unigenes that are generally taxonomically coherent. This rationale is conceptually analogous to the construction of metagenome-assembled genomes^58^ (MAGs).

To generate MGT v1.5, we rebuilt the collection from scratch, incorporating the Tara Polar Circle samples. Briefly, an abundance matrix was computed using RPKM (reads per kilobase per million mapped reads) for unigenes detected frequently enough across samples. This matrix was then submitted to the canopy clustering algorithm described by Nielsen *et al*.^103^, a density-based method that relies solely on abundance covariation. We used a max Pearson’s correlation difference of 0.1 to define clusters and then clusters were merged if canopy centroids’ distances were smaller than 0.05 (250k iterations, default parameters). This updated version extends the collection to a total of 1000 MGTs counting at least 1000 unigenes each (previously 924 MGT with at least 500 unigenes, +8.2%) encompassing 9,547,691 unigenes (previously 6,946,068 +37.5%).

### Environmental parameters and distance

We used a set of nine environmental parameters from four different sources (*In situ*, World Ocean Atlas 2018, PISCES-v2 model and the U.S. Naval Observatory), all believed to be relevant ecologically. We used three *physical* variables: Temperature and Salinity (*in situ Tara* measurements^105^) and Sea Sunshine Duration (SSD, Calculated from Astronomical Applications of the U.S. Naval Observatory). Two macronutrients were taken from *in situ* measurements: dissolved Silica (Si0^2^), and phosphate (PO_4_^3-^)^106^. Nitrate 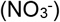 was extracted from the World Ocean Atlas 2018^107^.

In addition, two seasonality indices (for temperature and nitrate) were calculated using data from the World Ocean Atlas 2018 to reflect seasonal maximal seasonal fluctuation over the years at the sampling locations:

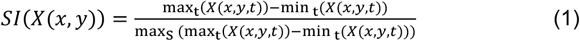

*X* is either nitrate concentration or temperature. The indices *t* and *S* respectively refer to time (months here) and spatial coordinates. All variables are extracted at the month and sampling locations considered here.

An environmental distance was calculated as the Euclidean distance between stations using z-score standardized values of each parameter):

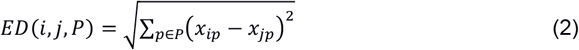

*ED*(*i, j, P*) is the environmental distance between stations i and j for a set of parameters P (here we used the whole set).

### Remotely sensed environmental variables

To visualize the context of sea surface temperature around the polar front and at the sampling date, we used re-analyzed remotely sensed Sea Surface Temperatures data from NOAA OI SST V2 High Resolution Dataset corresponding to June 2013^108^. For Chlorophyll *a* and Net Primary Production, we used remotely sensed estimates for June 2013 from an ocean color algorithm developed for the AO^41,42^.

### Identification of the polar front

To determine the position of the polar front from the SST estimates, we adapted the methods from Oziel *et al*.^99^ and Neukermans *et al*.^27^. These studies define the polar front as a transition zone mostly situated within 0°C and 3°C winter (March-April) surface isotherms. Atlantic Waters are usually defined as waters warmer than 3°C and Arctic Waters as water colder than 0°C. Isotherms of 0°C and 3°C of the SST product are plotted using the ggplot2^109^ geom_contour function in R.

### Calculating -omic community dissimilarity and the atlanticity indexes

Following Richter *et al*.^40^, we estimated pairwise community dissimilarity using shotgun metagenomes and metatranscriptomes from the size fraction 0.8-2000 µm. To do so, we used SimkaMin^100^ rather than Simka^100^, an optimized method for comparing -omic datasets based on k-mer frequencies (short DNA subsequences of length k). It is also more computationally efficient due to a subsampling-based approximation scheme that reduces memory usage and runtime without compromising accuracy to estimate Bray-Curtis and Jaccard dissimilarities. *β diversity* for -omic reads is defined with the following equation within Simka:

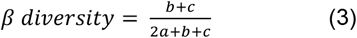

Where a is the number of distinct k-mers shared between two samples, and b and c are the number of distinct k-mers specific to each sample. We represented the distance between each pair of samples on a heatmap using the heatmap.2 function of the R-package gplots_2.17.0^104^. We also calculated β-diversity between stations *i* and *j* based on MATOU-v2 unigenes abundances as follow:

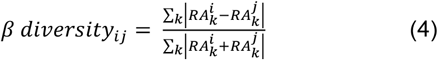

Where 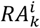 is the relative abundance of unigenes *k* in station *i*. We obtained high correlation between unigenes based distance and simkaMin based distance (r^2^=0.96 for metatranscriptomes-based distances) as well as between metatranscriptomes and metagenomes distances (r^2^=0.99 and 0.97 respectively for simkaMin and unigenes Bray-Curtis).

Using either measure of β*D*, we calculated a single metric of “atlanticity”, named here as the atlanticity genomic index. It is calculated as follows:

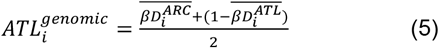

Where 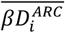 and 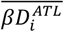 are the average *βD* value of station *i* within the AO, respectively the NAO cluster. The index has values between 0 and 1 and is symmetric with the “Arcticity” index: *ARC*_*i*_ = 1 − *ATL*_*i*_.

Second, for each unigene, we define the atlanticity geographic index based on presence/absence of genes across AO and NAO station as follows:

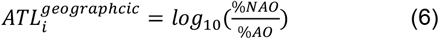

Where %*NAO*and %*AO* are the percentage of NAO, respectively AO samples in which the unigene is present. The index is comprised between -2 and 2 and rescaled between -1 (fully AO unigene) and 1 (fully NAO unigene).

### Taxonomic and geographical annotations of unigenes

Each sample of our dataset is characterized by the taxonomical annotation of unigenes. To harmonize taxonomic annotation, we focused on a subset of sufficiently abundant groups in our dataset: Hexanauplia, Bacillariophyta, Pelagophyceae, Phaeocystales, Ciliophora, Tunicata, Mamiellales, Dinophyceae, Rhizaria, Unclassified Eukaryota, as well as other Haptophyta, Opisthokonta, and Stramenopiles. Unigenes not assigned to any of these groups were annotated as ‘other’. This category included Streptophyta, Cnidaria, Rhizaria, Insecta, Cryptophyta, Euglenozoa, Amoebozoa, Craniata, Fungi, Viruses, Coccosphaerales, Archaea, root and other Alveolata. Each unigene was also assigned into one of three types of geographical annotation based on its distribution: present only in the NAO, present only in the AO, or shared between the two basins (detected in at least one station of each basin). To identify taxonomic groups or clades that are significantly more relatively abundant in one basin compared to the other, we applied a Wilcoxon test on the mean of the log-transformed relative abundances of each taxon across stations of the two basins. We also applied a Hommel correction^110^ to adjust *p*-values for multiple hypothesis testing. Note that for this analysis we used the log-transformed values of relative abundances rather than centered log ratio (which was used for the unigenes expression analysis, see below) as CLR might distort interpretation (as it is centered on zero) in the case of skewed distributions (which is the case here).

### Functional characterization: comparison between AO and NAO

For each unigene from the MATOU-v1.5 catalog, we converted PFAM^111^ (protein family domain) annotation to corresponding Gene Ontology (GO) terms^112^. We used the full GO dataset including the three main categories of the catalog: molecular function, cellular component, and biological processes. Since a single unigene can contain multiple PFAMs and a single PFAM can correspond to multiple GO terms, we expanded the functional annotations accordingly. Each unigene was assigned to every GO term within its associated GO hierarchy, possibly resulting in multiple functional annotations per unigene. For each GO term found across our dataset (a total of 4,695 terms out of a total of 44,000 in the database), we defined its relative abundance per sample by summing the relative abundances of all unigenes annotated with that term.

To detect functional differences between the Arctic Ocean (AO) and North Atlantic Ocean (NAO), we applied Student’s t-tests to compare the means of the log-transformed relative abundances of each GO term across samples from each basin. We performed two types of multiple testing correction: Hommel correction^110^, and False Discovery Rate (FDR) correction^113^ at a threshold of FDR = 0.05.

The two methods yielded different numbers of significantly more abundant functions (see Main Text), the Hommel method being more stringent. Here we used only the log-transformed relative abundances rather than centered-log-ratio. The multiplicity and unevenness of levels in the GO hierarchy didn’t allow us to define a robust mean. The same statistical test was also conducted to select taxonomic groups of interest independently.

To identify significantly overabundant PFAMs between NAO and AO metagenomes, we used ALDEx2 R package^53,54^. ALDEx2 uses standard Bayesian inference^114^ to estimate the posterior distribution of [*p*_1_, *p*_2_, …, *p*_*i*_, …, *p*_*n*_] as the product of a multinomial likelihood (where *p*_*i*_ is the proportions of gene *i*, here PFAMs, in a sample with a 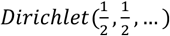 prior. ALDEx2 uses the full Dirichlet posterior probabilities distribution to draw multiple estimates (Monte-Carlo realizations) of each *p*_*i*_ such that 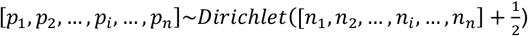. Drawing *p*_*i*_ from the full posterior allows accounting for the large variance and extreme non-normality of the marginal distributions of *p*_*i*_ when the associated *n*_*i*_ are small. *p*_*i*_ are then transformed using the centered-log-ratio (see the next paragraph for this transformation). ALDEx2 therefore models the abundance of each gene, here PFAMs, as a random-effect ANOVA model accounting for the fact that within-condition abundance/expression is a non-negligible random variable:

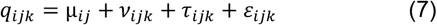

Where *q*_*ijk*_ is the adjusted log-abundance/expression, *μ*_*ij*_ is the expected expression of gene *i* within each condition *j, ν*_*ijk*_ is the sample specific expression change for replicate *k, τ*_*ijk*_ is the sampling variation from inferring abundance from read counts and *ε*_*ijk*_ the remaining nonspecific error, independent and identically distributed within *k*. To find significantly overabundant PFAMs, ALDEx2 then performs a Welch t-test on each *q*_*ijk*_ estimates (drawn from the Monte Carlo realization) and corrected for multiple comparison by the Benjamini-Hochberg method^115^.

### Community composition and expression-based distances

We computed pairwise community composition and expression distances between stations using centered log ratios (CLR) transformed PFAM abundances estimated from the ALDEx2 package^53,54^ Expression was defined as the difference between CLR-transformed metatranscriptomic abundances and the CLR-transformed metagenomic abundances (Equation 8). Then, the distance between each pair of stations was defined as the Euclidean distance over all PFAMs:

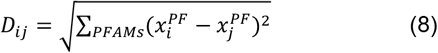

Where 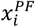 is either the CLR-transformed abundance or the expression value, as defined above, of PFAM *PF* in station_*i*. Following the approach described in Salazar *et al*.^8^, we then computed, for temperature bins containing 5 stations, the median ratio between abundance-based distances and expression-based distances of all pairwise distances among stations of each bin. To test whether this median ratio was significantly different than 1, a Wilcoxon test and holm correction for multiple comparison (p<0.05) were applied for each bin.

### Embedding of the gene expression matrix using PHATE

We defined the expression level of each unigene as the difference between its centered-log-ratio (CLR)-transformed relative abundance in metatranscriptomic reads and its CLR-transformed relative abundance in metagenomic reads:

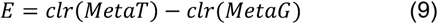

The CLR transformation was used as it was shown to be stable to subsampling of the dataset^116,117^. It was defined as follows:

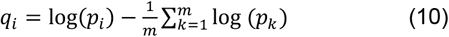

Where *p*_*i*_ was the relative abundance of unigene *i* and we have 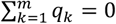.

We used the PHATE dimension reduction algorithm (Potential of Heat diffusion for Affinity-based Transition Embedding)^55^ on the gene expression matrix from approximately 572,000 unigenes that were present in both metatranscriptomic and metagenomic dataset across at least 5 stations. This threshold considerably decreased the number of analyzed unigenes but was chosen to ensure a most robust statistical power and to integrate both metatranscriptomic and metagenomic abundances.

To validate this approach, we computed for each unigene, the Erb rho correlation coefficient^118^ between *clr*(*MetaT*) and *clr*(*MetaG*). Importantly, these two measures correlated strongly with each other although depending on the taxa, supporting the robustness of this approach (Figure S10).

As PHATE cannot handle missing values, we set to zero (which is the mean expression value across all unigenes) the expression of unigenes for which we could not define an expression value in any given station. Although this imputation was not fully satisfying, it was the most effective way we found to preserve gene expression variation signals in our dataset. Alternatively setting to 0 the ‘real’ zeros and renormalizing expression values to positive values proved inadequate, as it distorted the signal in reason to many zeros. Therefore, the PHATE embedding may not fully capture correlations between gene expression and environmental parameters, as many expression values are set to 0. This limitation is also discussed in the section on correlation with environmental parameters.

Briefly, PHATE allowed the visualization of high-dimensional data while preserving both local and global structures. To generate the embedding, PHATE first transforms the Euclidean distance matrix using an adaptive, non-linear kernel. This transformation emphasizes local similarities and minimizes noise, allowing for the detection of local expression patterns within the data. After this initial step, PHATE captures the global structure of the dataset through a random-walk diffusion process by factoring the diffusion operator (the fitted kernel) to a certain optimized power. This operation allows capturing long-range branching events in the dataset and diminishing the local noise. Finally, PHATE computes the log-transformed diffusion probabilities and embeds them into a lower-dimensional space (we used a two-dimensional embedding here) using metric multidimensional scaling MDS.

One important hyperparameter in the PHATE algorithm is *k*-nearest neighbor (*kNN*) that determines the bandwidth of the kernel during the first step of the embedding process. For each data point, this bandwidth is set to the distance to its *k*-th nearest neighbor, thereby allowing the kernel to adapt locally to the density and structure of the data. Given the large size of our dataset (572,000 unigenes), we set *k NN* to 500.

### Metabolism annotation using PFAM domains

We annotated metabolic functions related to photosynthesis, including components of photosystems, Light Harvesting Complexes (LHCs), and Cyclic Electron Transfer (CET) using the Pfam protein family domain database^111^. Annotation was performed by mapping PFAM domains to unigenes in our dataset. The full list of PFAM domains used for this analysis is provided in Table S3.

### Correlation of unigenes expression with environmental parameters

To assess the influence of environmental conditions on gene expression, we retained only unigenes that were detected in at least five stations and present in both basins (those with at least one expression value in both AO and NAO). This filtering reduced the unigene set from approximately 572,000 to approximately 485,000 unigenes.

For each unigene expression profile, we calculated two Pearson correlations coefficient with each environmental parameter: one using the actual expression values and one another using shuffled expression values as a null model. A unigene was considered significantly correlated with a given environmental parameter if its observed correlation value fell within the extreme 5% of the null distribution (i.e., below the 2.5th or above the 97.5th percentile). This corresponds to a false discovery rate (FDR^113^) of 0.05.

Then, to further characterize the functional landscape of unigenes significantly correlated with environmental variables, we performed GO term enrichment analyses. These were based on the deepest level of GO annotation associated with each unigene. A hypergeometric test was applied to assess overrepresentation, using the full set of unigenes from the same taxonomic group or MGT as background. *P*-values were corrected for multiple testing using the Hochberg procedure, with a significance threshold of *p* < 0.05^119^. All analyses were conducted separately for each taxonomic group and each metagenomics-based transcriptome (MGT).

## Notes

https://www.genoscope.cns.fr/tara/

https://github.com/PaulFremont3/metaT_AO_NAO

https://doi.org/10.5281/zenodo.17316416

https://doi.pangaea.de/10.1594/PANGAEA.875582

https://psl.noaa.gov/data/gridded/data.noaa.oisst.v2.highres.html

http://albedo.stanford.edu/gertvd/research/arctic/prod/

https://www.genoscope.cns.fr/tara/

### Competing Interest Statement

The authors have declared no competing interest.

### Summary of Updates

Updated abstract; Updated Figure 1, Figure S8, Table S1; Updated Data availability statement;

