## Supplementary_Information for "Changes in gene expression in eukaryotic phytoplankton at the Atlantic-Arctic polar front"

Frémont *et al.*

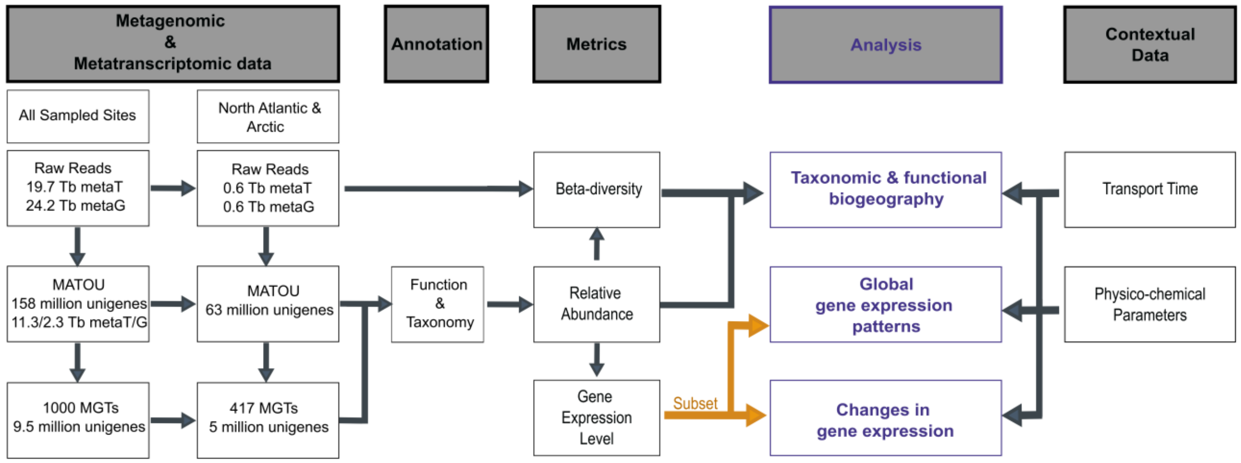

**Figure S1. Pipeline of the study.**

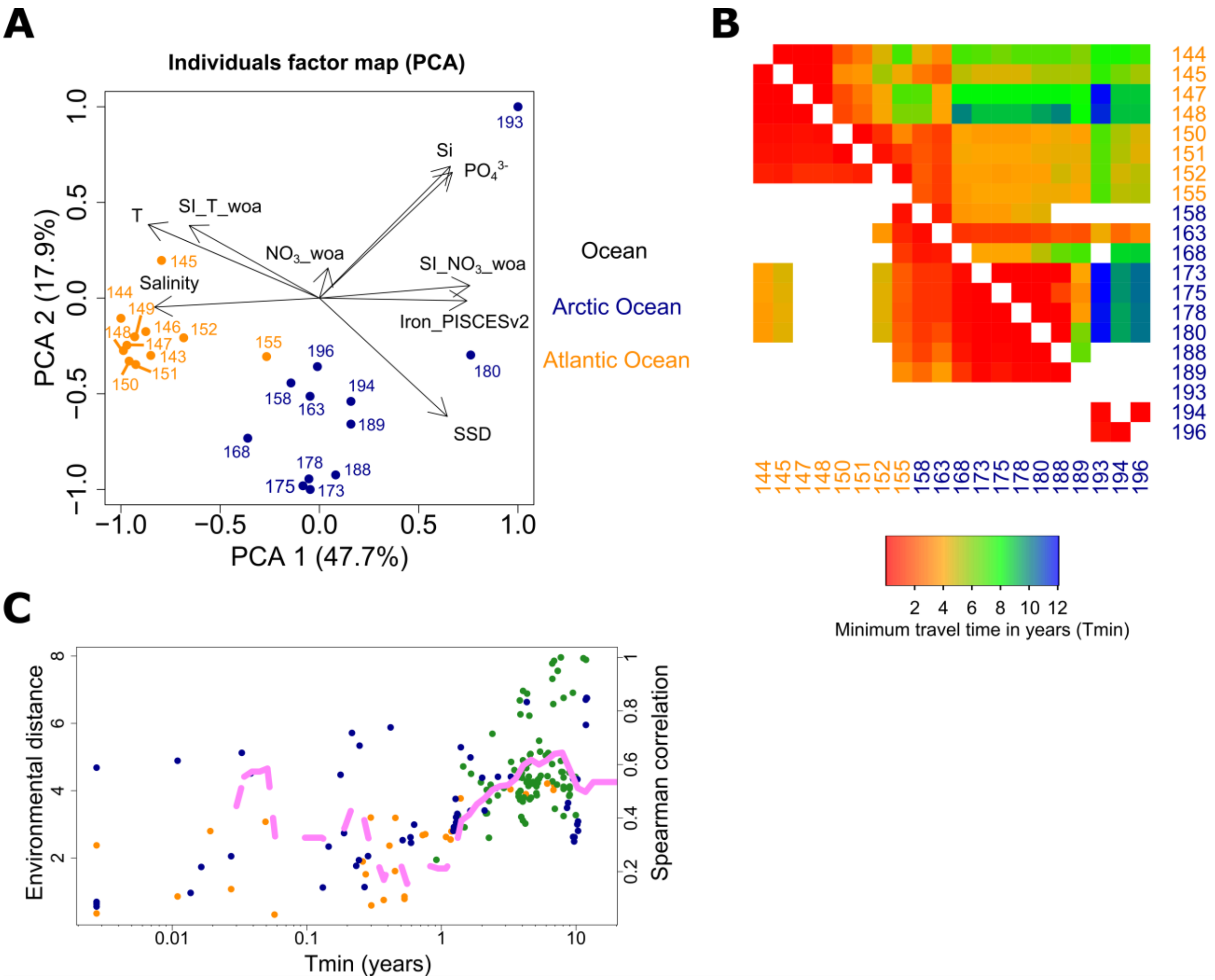

**Figure S2. Environmental differentiation of the NAO and AO basins with respect to transport time by currents.** (**A**) Principal Component Analysis (PCA) of the environmental parameters characterizing the sampled stations. (**B**) Minimum travel time connectivity matrix. The upper matrix represents the minimum travel time (Tmin) by ocean currents from row stations to column stations and vice versa. (**C**) Environmental distance in function of minimum travel time (Tmin) and cumulative Spearman correlation (pink line). When the line is full, the correlation is significant ($p<0.05$).

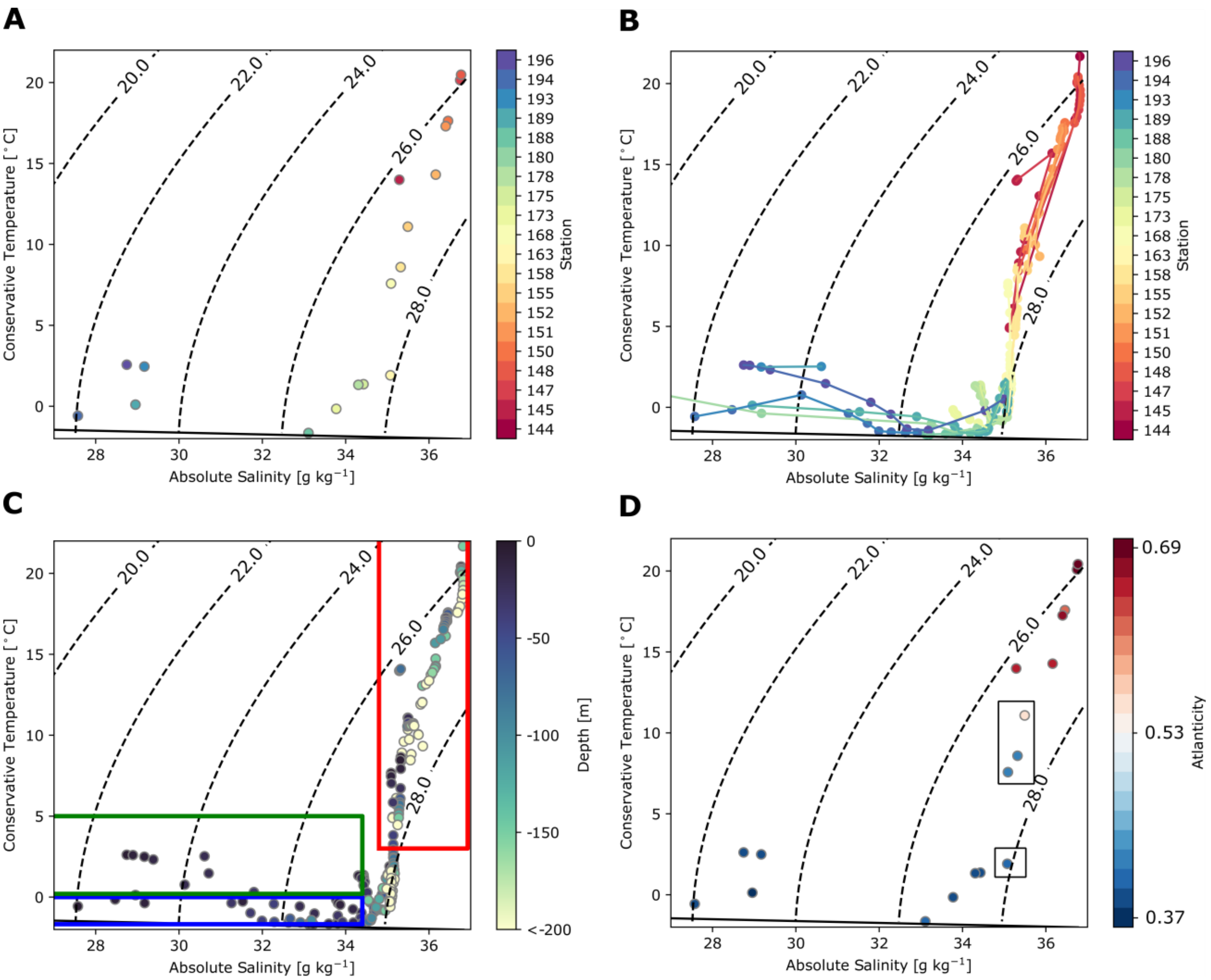

**Figure S3. Water mass determination.** Conservative temperature (°C) versus absolute salinity ($g.kg^{-1}$) diagrams for all stations sampled with (**A**) sample values and corresponding station identification number, (**B**) Full CTD profiles and point colored according to the station number, (**C**) full CTD profiles and point colored according to depth. The blue box defines the Arctic Waters (Tc < 0°C and Sa < 34.4), the red box marks the Atlantic Waters (Tc < 3°C and Sa > 34.7) and the green box illustrates the melt and coastal surface waters (Tc > 0°C and Sa < 34.4). Points at the bottom right corner correspond to modified Atlantic Waters. (**D**) Surface values and the atlanticity genomic index based on genomic differences between stations. Stations in the black frames are transitions stations near the polar front: 155, 158 and 163 and 168. Water masses definitions were adapted from Oziel *et al.* [S1]. The black line represents the freezing point of water at the surface.

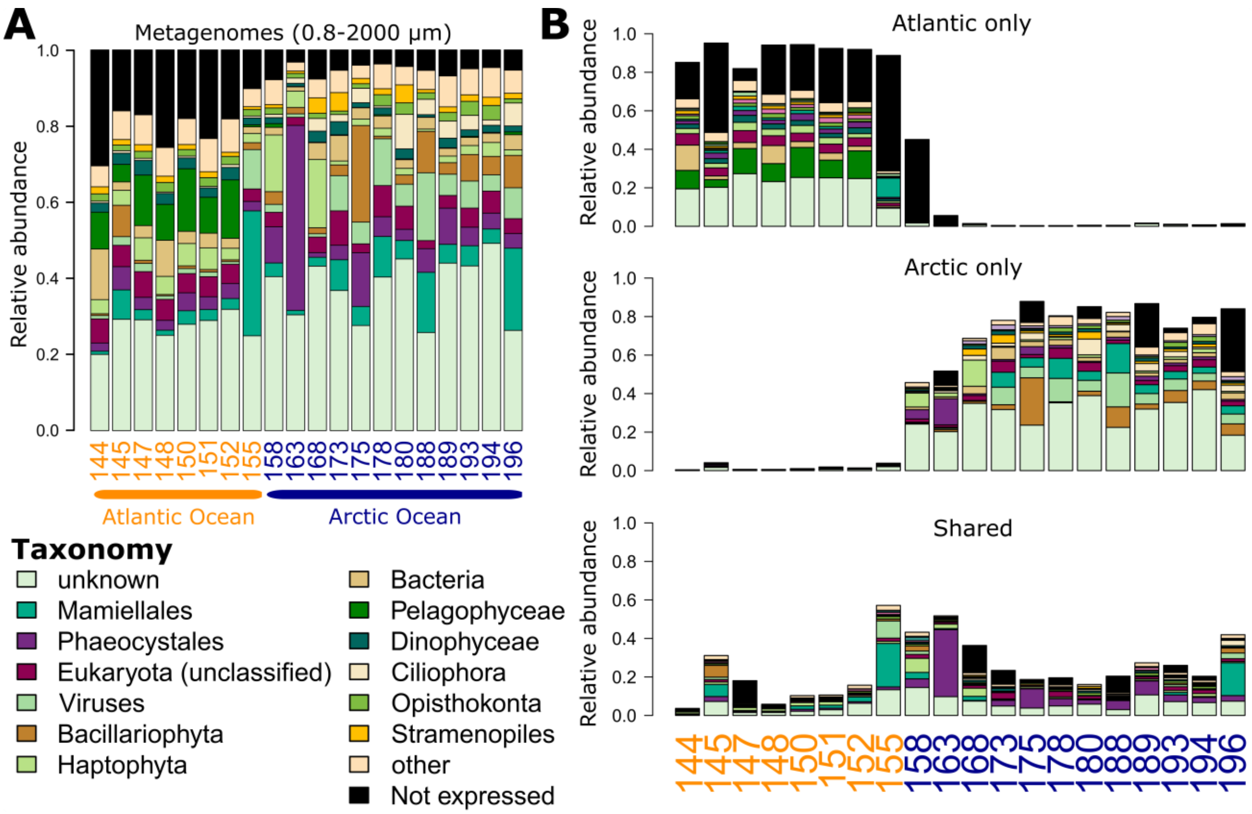

**Figure S4. Taxonomic composition of surface ocean plankton metagenomes.** (**A**) Global metagenomic taxonomic composition based on expressed unigenes relative abundances of major plankton taxa (taxonomic level can vary) in the NAO and AO. (**B**) Decomposition of basin-specific unigenes and shared unigenes between the two basins, based on presence or absence of expression. A marked transition enriched in shared unigenes appears at stations 155, 158 and 163. Data shown are for the 0.8-2000 µm size fraction only.

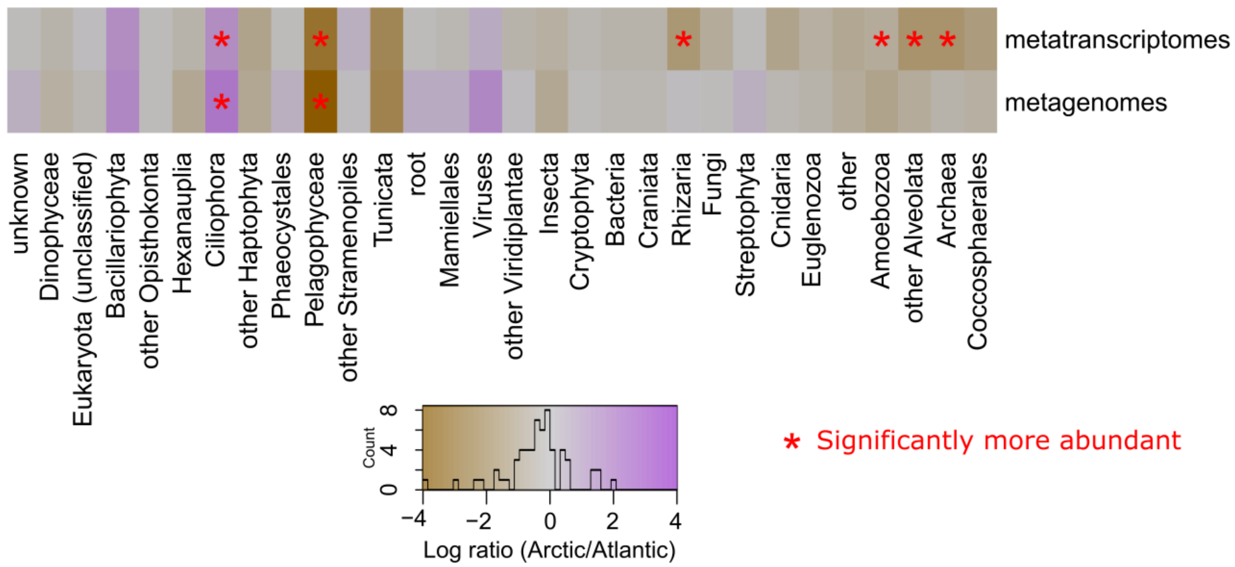

**Figure S5. Median log$_e$-fold change in relative abundance between Arctic Ocean (AO) and North Atlantic Ocean (NAO) metatranscriptomes and metagenomes for main plankton groups.** The heatmap represents, for each taxon, the difference in median log-transformed relative abundance between the two basins. Taxa significantly more abundant ($p<0.05$, Wilcoxon test with Hommel correction) in metatranscriptomes (top row) or metagenomes (bottom row) are indicated with red stars.

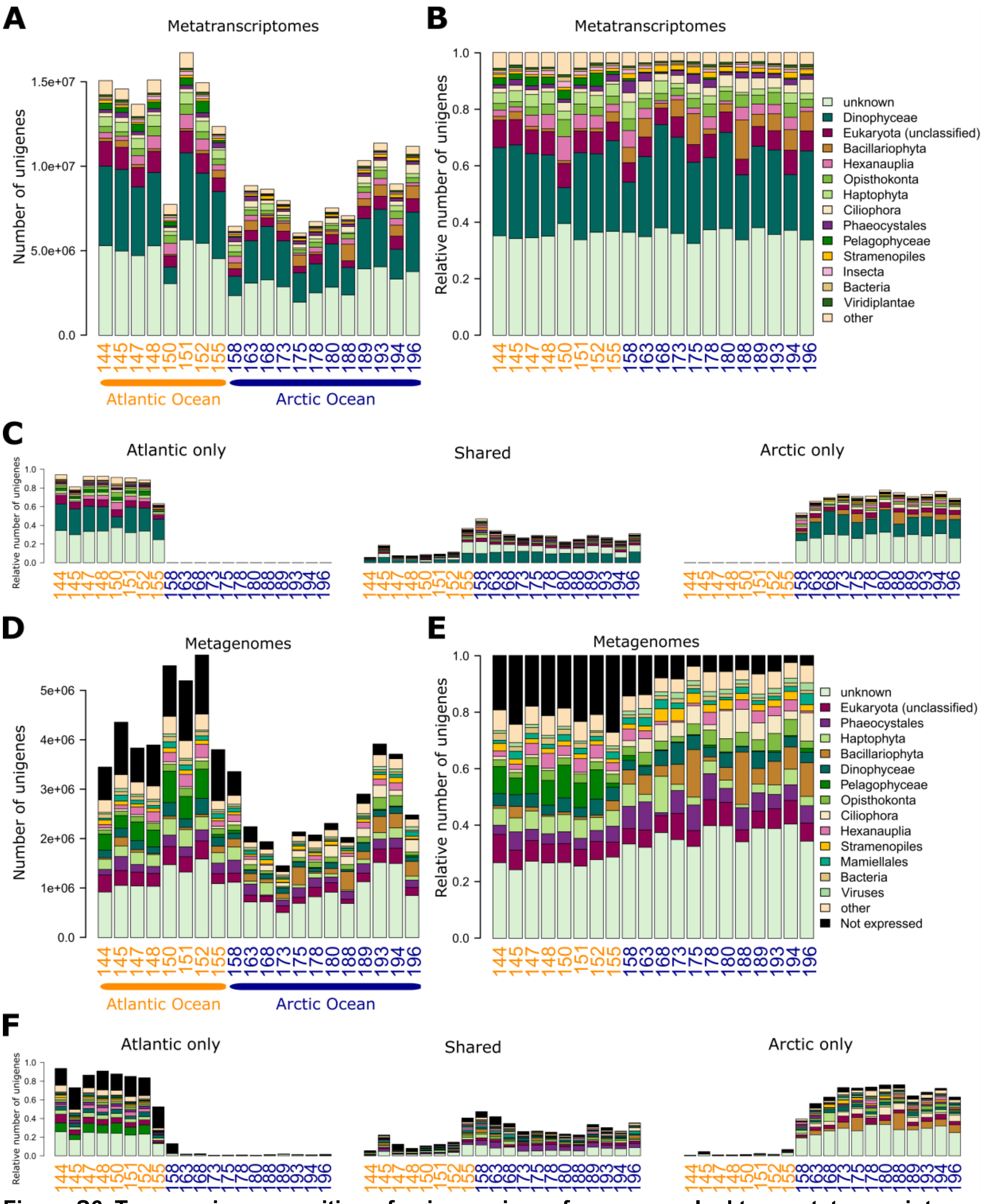

**Figure S6. Taxonomic composition of unigenes in surface ocean plankton metatranscriptomes and metagenomes.** (**A**) Metatranscriptomic and (**D**) metagenomic taxonomic composition based on number of unigenes at NAO and AO sampling stations. (**B**) Metatranscriptomic and (**E**) metagenomic taxonomic composition based on relative number of unigenes at NAO and AO sampling stations. (**C**) Decomposition of metatranscriptomes and (**F**) metagenomes based on relative number of basin-specific and shared unigenes between the two basins (classified based on presence or absence of expression). Data shown are for the 0.8-2000 µm size fraction only.

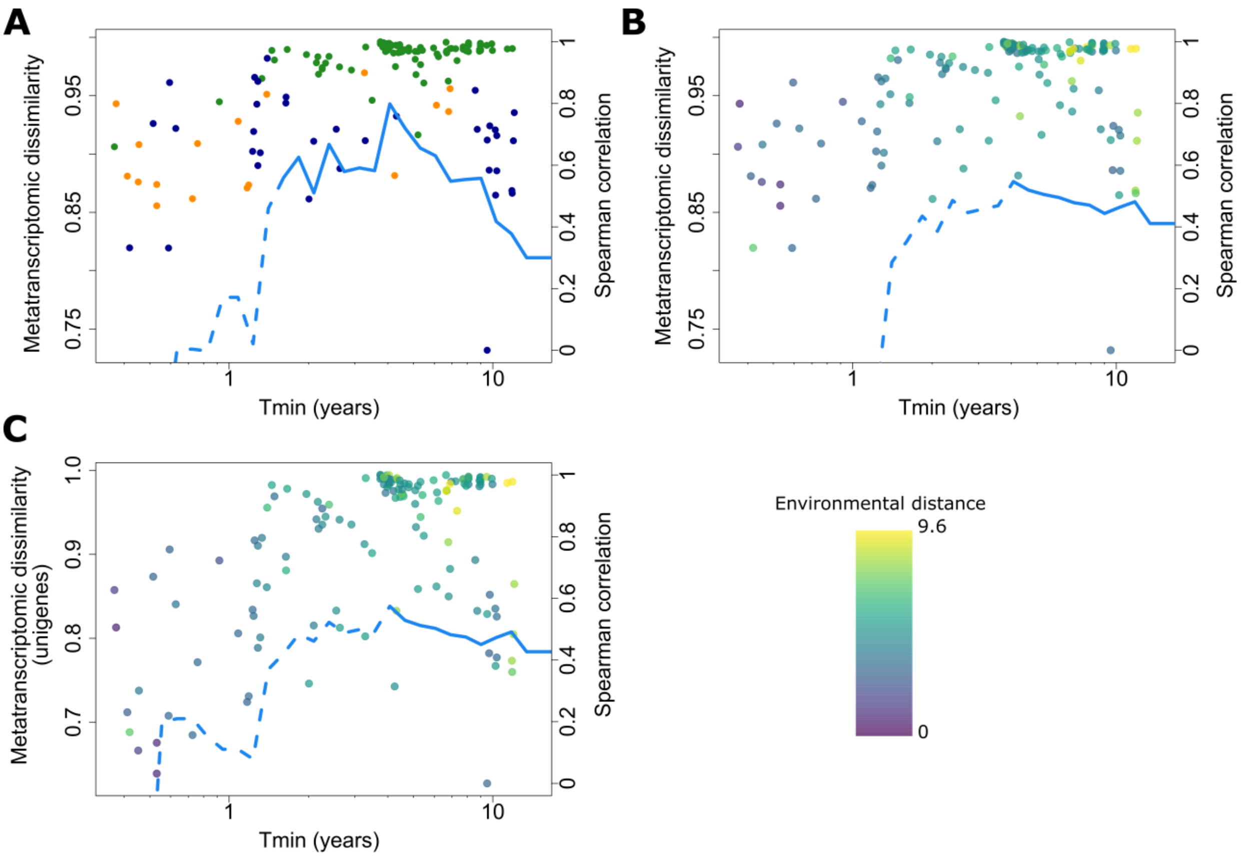

**Figure S7. Metatranscriptomic β diversity in relation to ocean current transport time and environmental distance between NAO and AO basins.** (**A**) Metatranscriptomic β diversity (simkaMin [S2]) in function of minimum transport time by currents between pairs of sampled stations and cumulative spearman correlation coefficient between the two metrics (blue line). Metatranscriptomic β diversity based on (**B**) simkaMin and (**C**) unigenes metatranscriptomic abundance-based Bray-Curtis in function of minimum transport time by currents between pairs of Tara station and cumulative spearman correlation coefficient (blue line) between the environmental distance (points' color) and β diversity.

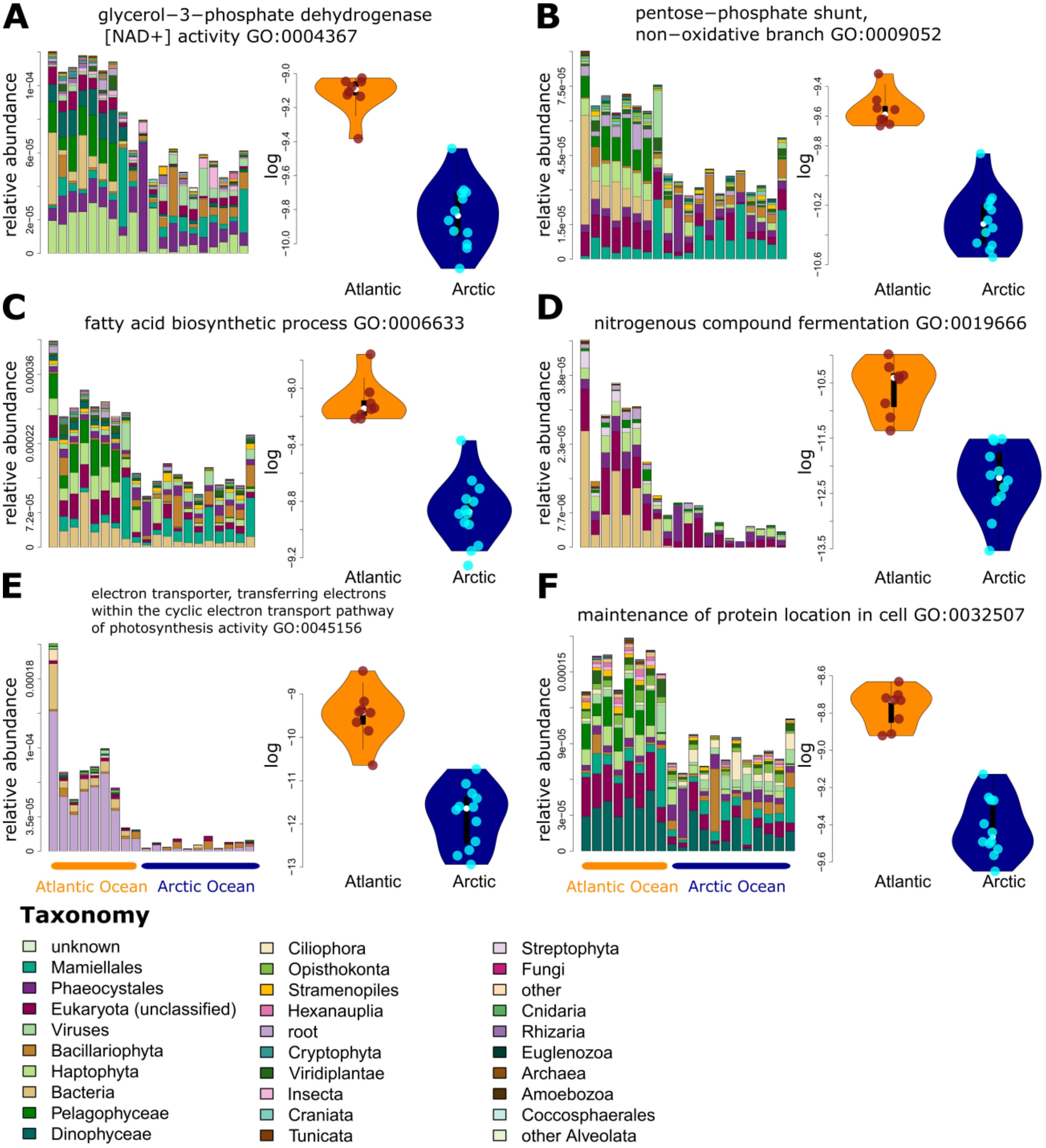

**Figure S8. Examples of Gene Ontology (GO) terms more abundant in the NAO metagenomes.** Each panel (**A**-**F**) shows, on the left, the total metagenomic relative abundance of unigenes associated with the abovementioned GO term across considered taxa, and on the right, the distribution of this relative abundance in the NAO basin (orange) versus AO basin (dark blue). Each of these GO terms was significantly more abundant in the NAO basin.

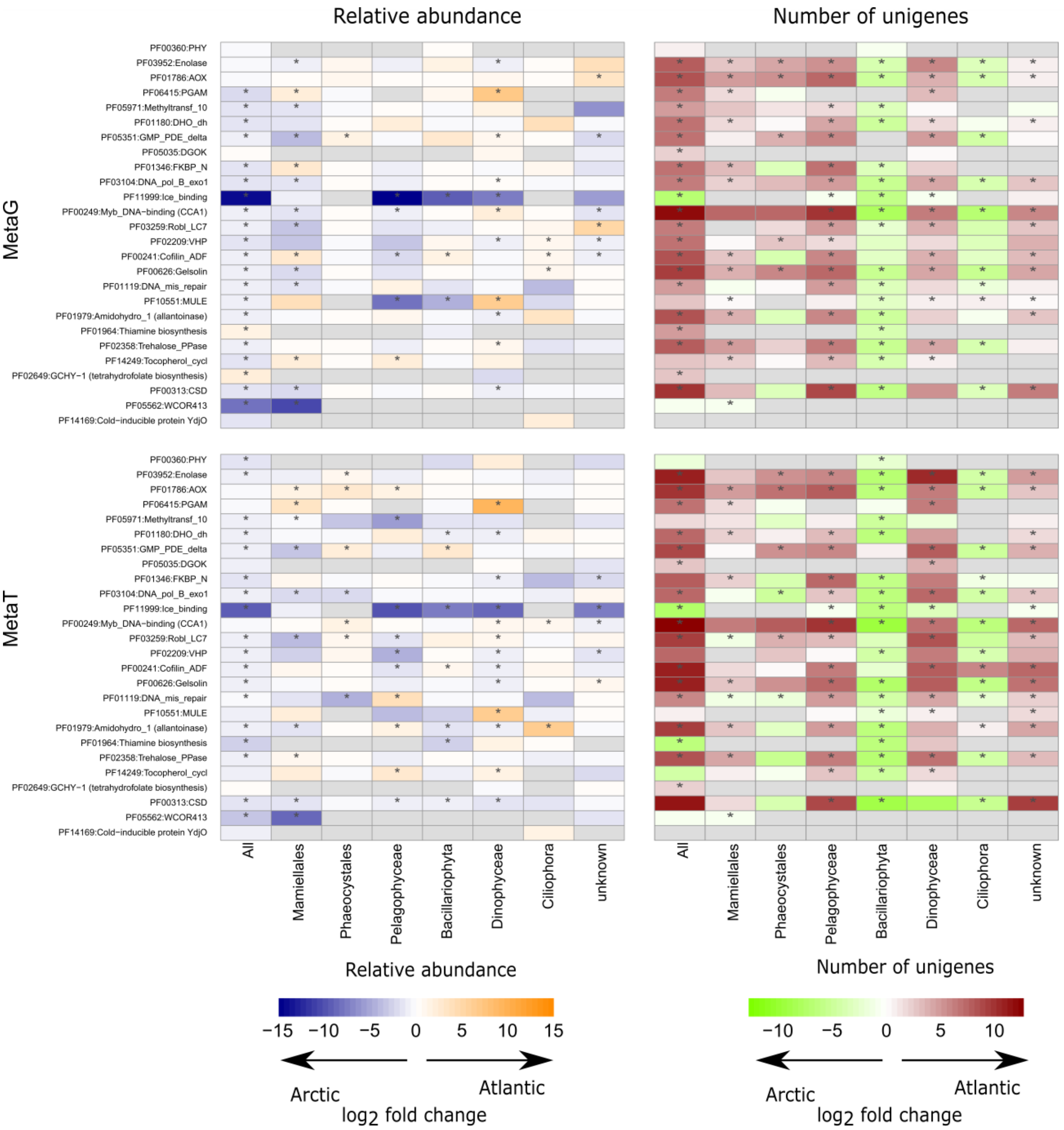

**Figure S9. Differential abundance of PFAMs associated to temperature response in metatranscriptomes and metagenomes across taxa.** Right column: Median $log_2$-fold change between NAO and AO metagenomes and metatranscriptomes for PFAMs associated to temperature response, based on estimated centered log ratios from the ALDEx2 package [S3,S4] for all taxa, Mamiellales, Phaeocystales, Pelagophyceae, Bacillariophyta, Dinophyceae, Ciliophora and unknown groups. Significant differences are highlighted by stars. Left column: Median $log_2$-fold change in the number of unigenes between NAO and AO metagenomes and metatranscriptomes for the different taxa.

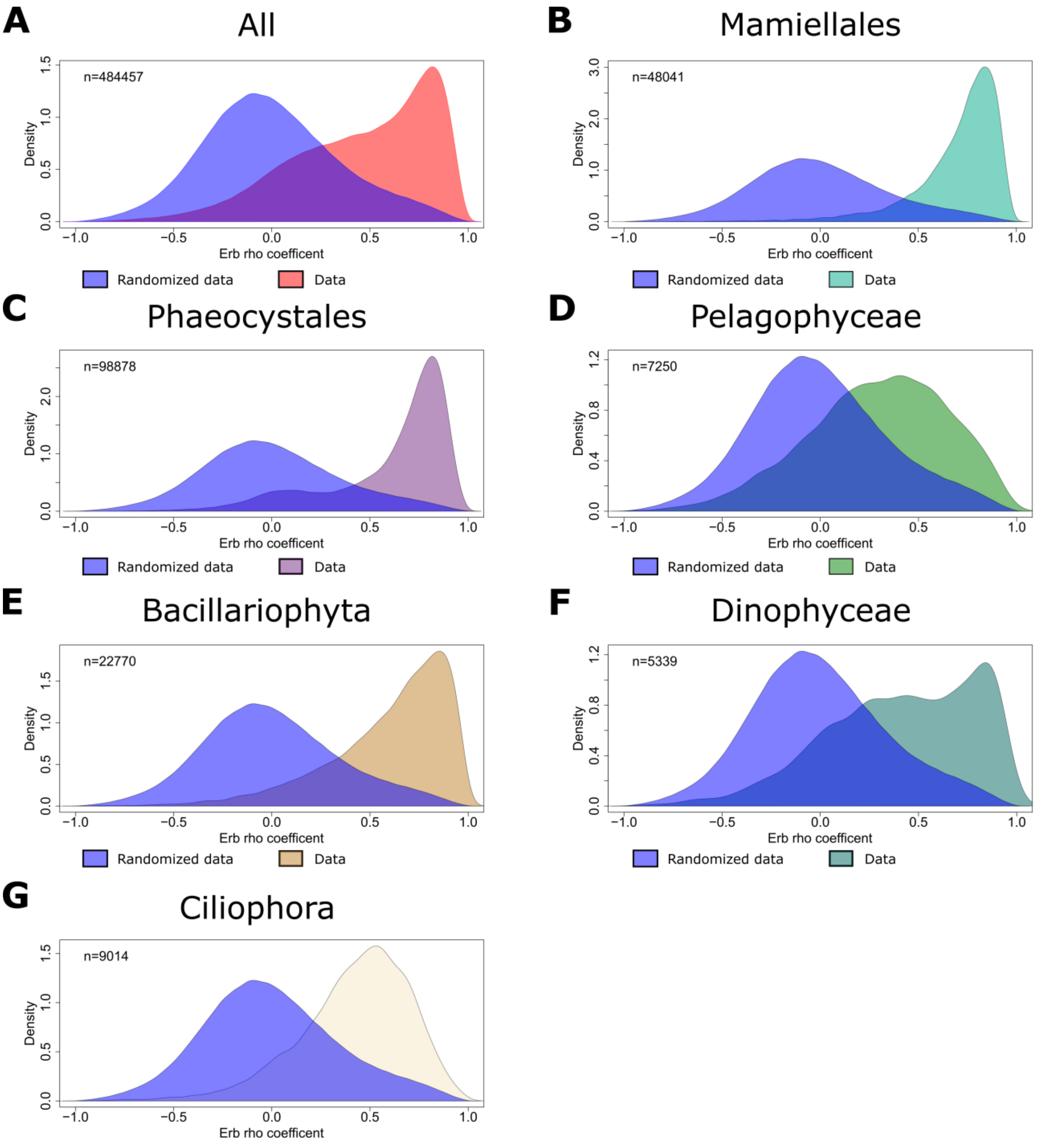

**Figure S10. Distribution of Erb's rho correlation index between centered log ratios of randomized and real metatranscriptomic and metagenomic relative abundance of unigenes present in at least five stations.** (**A**) Correlation index distributions for all taxa. (**B**-**G**) Distributions for individual taxonomic groups. Blue curves represent correlation distribution from randomized data; other colors correspond to real data. Importantly, metatranscriptomic and metagenomic centered log ratios (CLR) showed overall good correlation, with strength varying by taxon. These good correlations support the use of metagenomic CLR to normalize metatranscriptomic CLR in defining unigene expression levels.

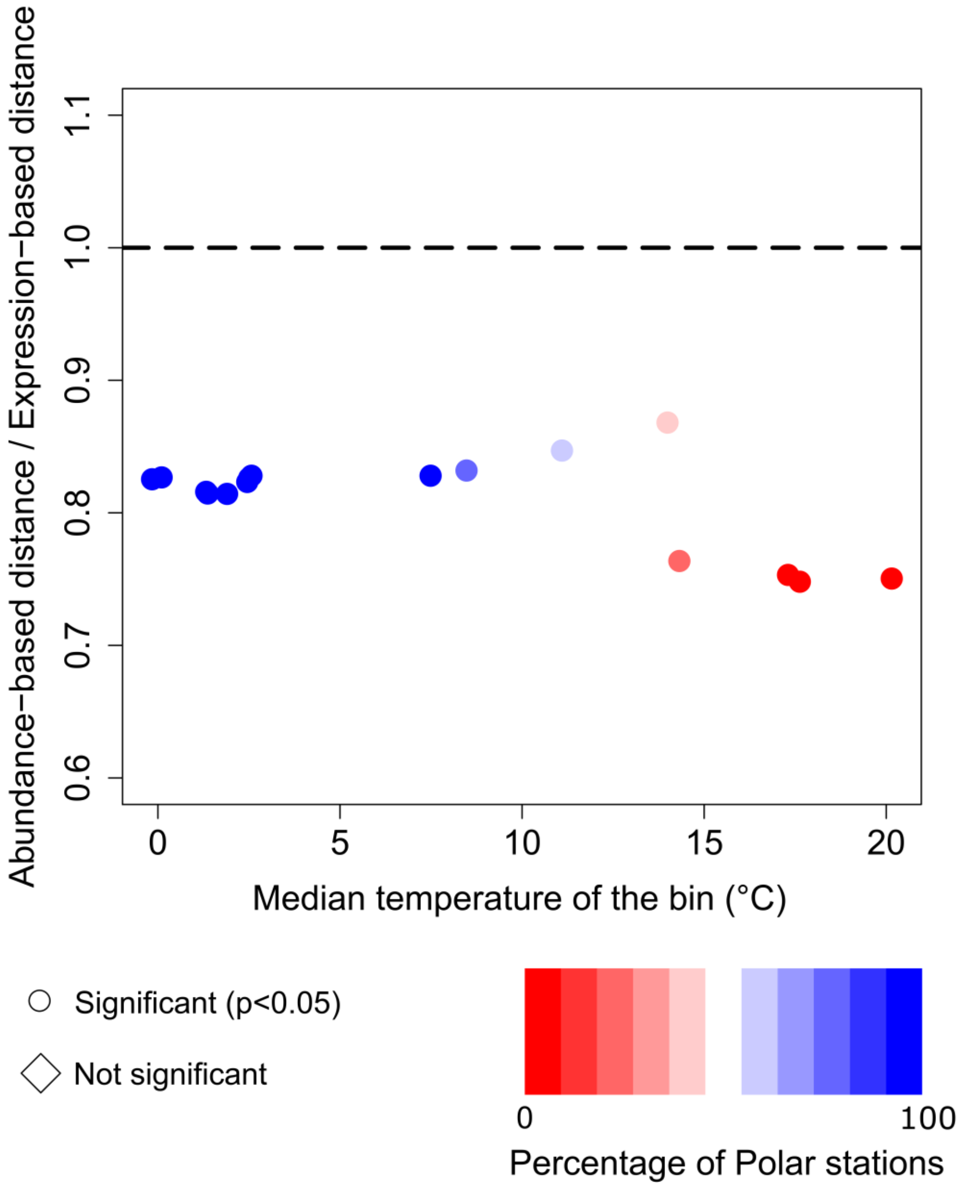

**Figure S11. Relative contribution of community composition and gene expression changes to variations in metatranscriptomic composition along the NAO-AO temperature gradient.** The plot shows the median ratio between an abundance-based distance and an expression-based distance (see Methods), calculated for temperature bins containing five stations each. A Wilcoxon test with Holm correction (p<0.05) was performed for each bin to test whether the median ratio significantly differed from 1.

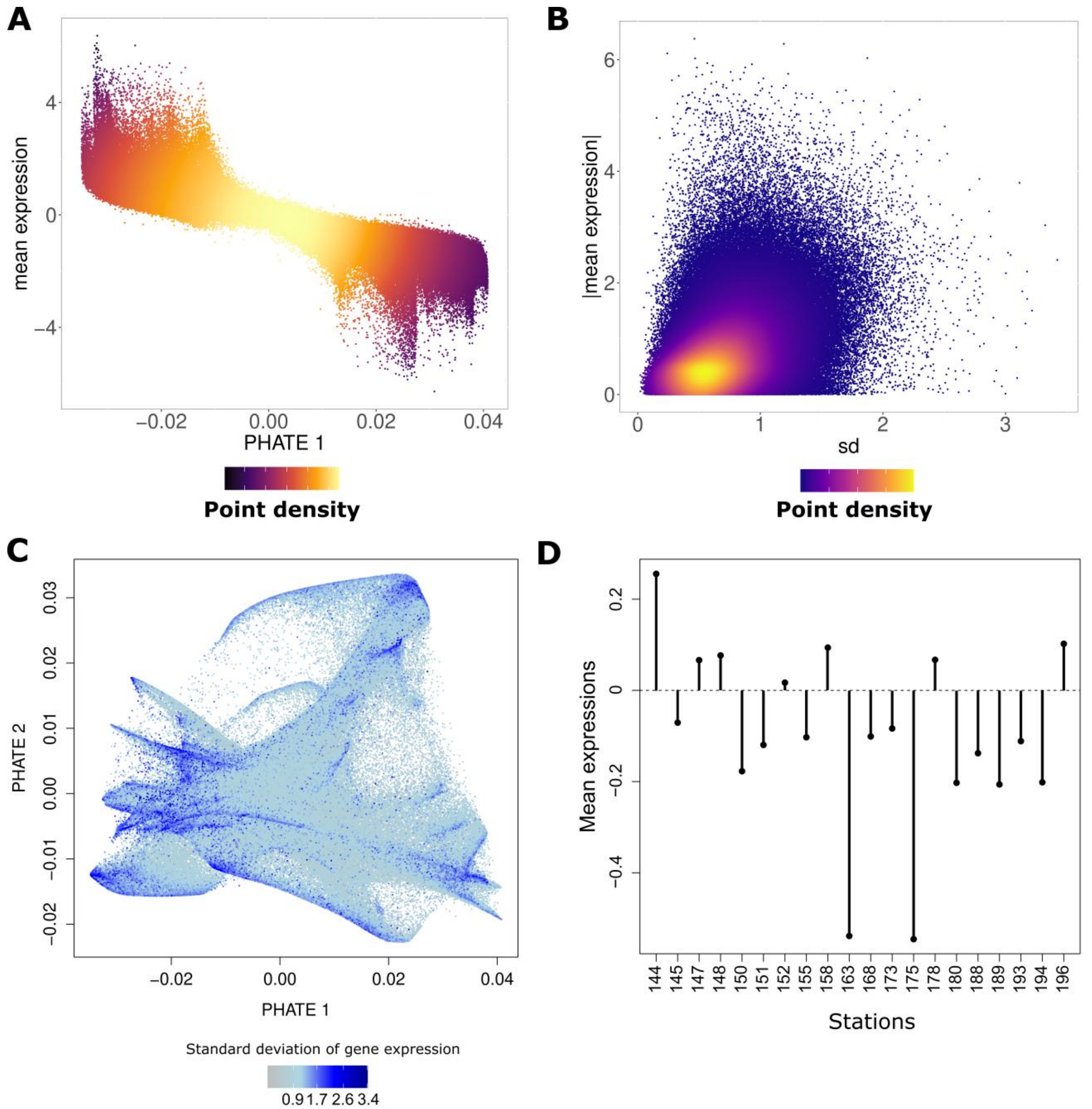

**Figure S12. Additional features explaining the PHATE structure.** (**A**) Mean expression versus PHATE 1 coordinates. (**B**) Absolute unigene expression value versus standard deviation of gene expression. (**C**) PHATE embedding with each dot colored by the standard deviation of gene expression. (**D**) Mean expression of the subset of the 572,000 unigenes used in the PHATE analysis as compared to 0 which is the mean expression in each station (CLR).

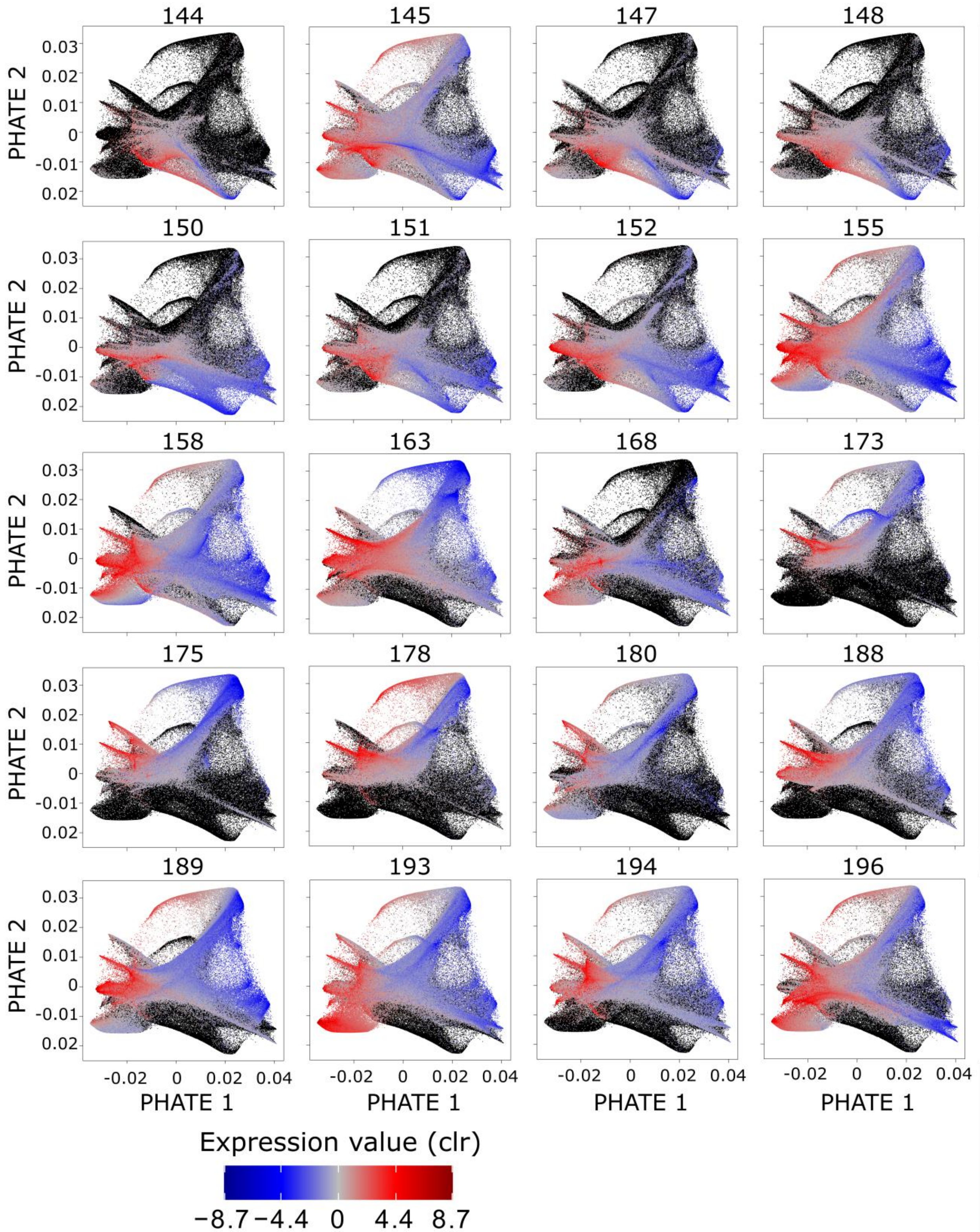

**Figure S13. Representation of individual sampled stations in the PHATE embedding.** For each station, genes are represented according to their PHATE coordinates. Unigenes are colored by their expression or represented in black when not expressed.

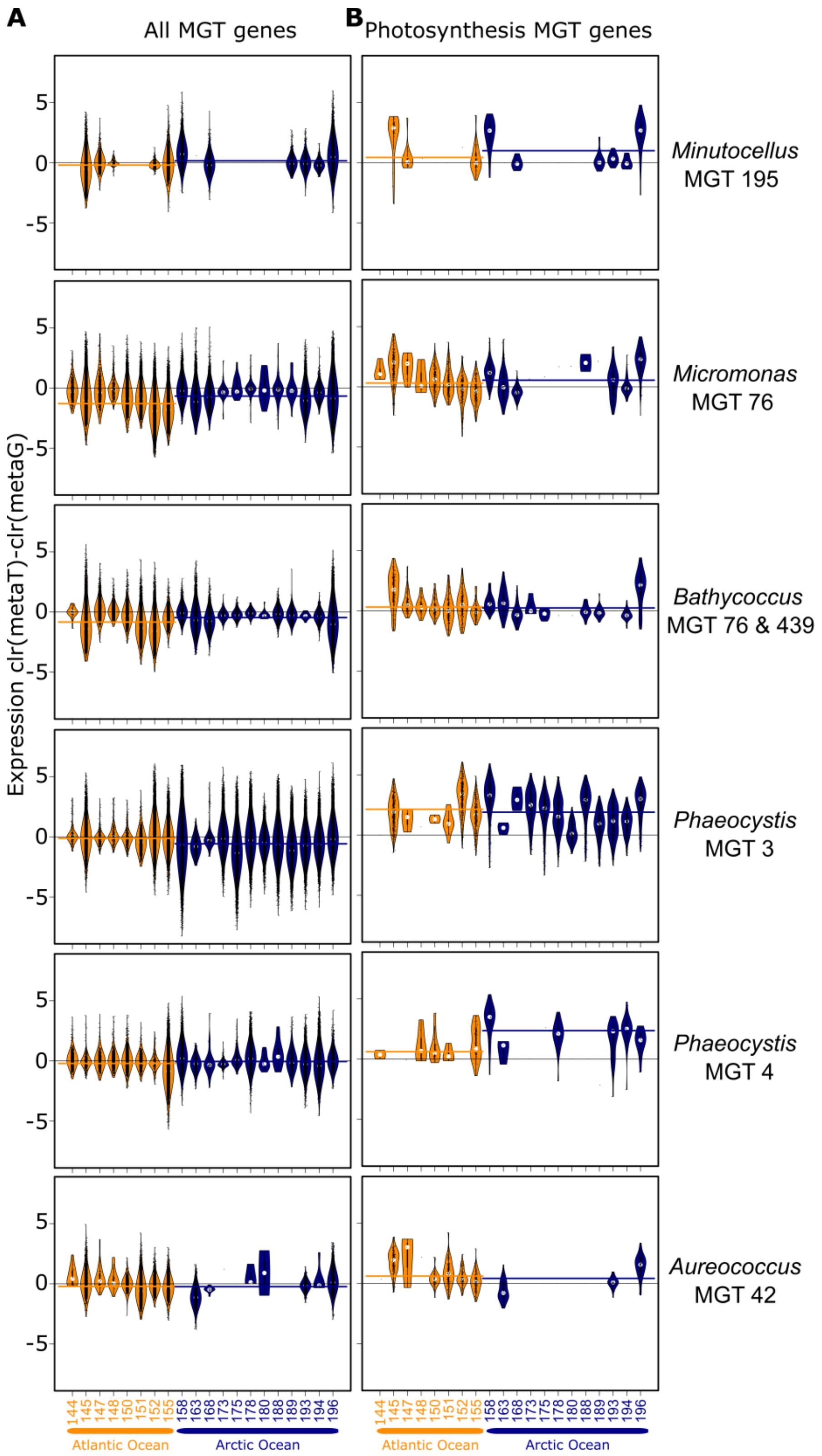

**Figure S14. Violin plots of gene expression across sampled stations for six metagenomics-based transcriptomes (MGTs).** (**A**) All genes. (**B**) Photosynthesis-related genes. Bars indicate the overall median expression per basin: orange for NAO and dark blue for AO.

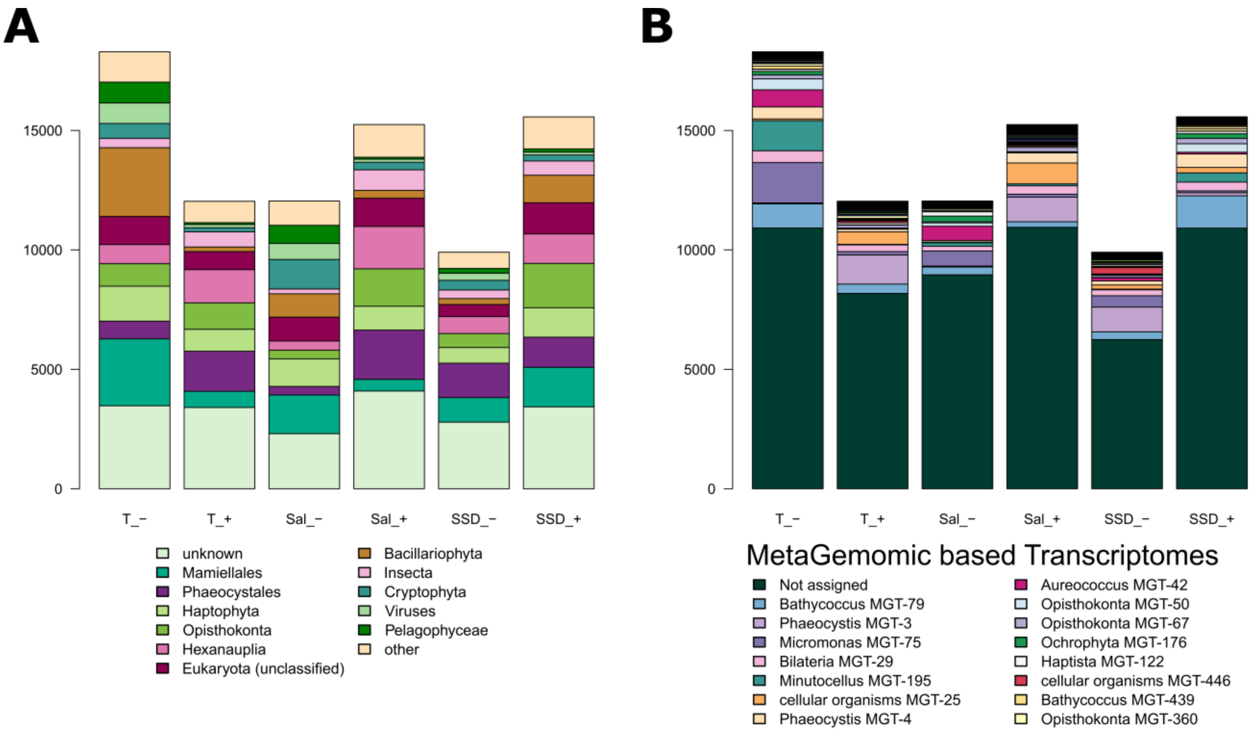

**Figure S15. Taxonomic decomposition of unigenes significantly correlated with physical environmental parameters, either negatively or positively.** (**A**) Taxonomic groups and (**B**) Metagenomics-based transcriptomes (MGTs) annotation of unigenes. Temperature is the tested parameter with the highest number of significantly correlated unigenes expression profiles. Because salinity (Sal) and day length (SSD) are highly correlated with temperature, we only analyzed unigenes functions whose expression is significantly correlated with temperature (Figure 7, S16 and S17).

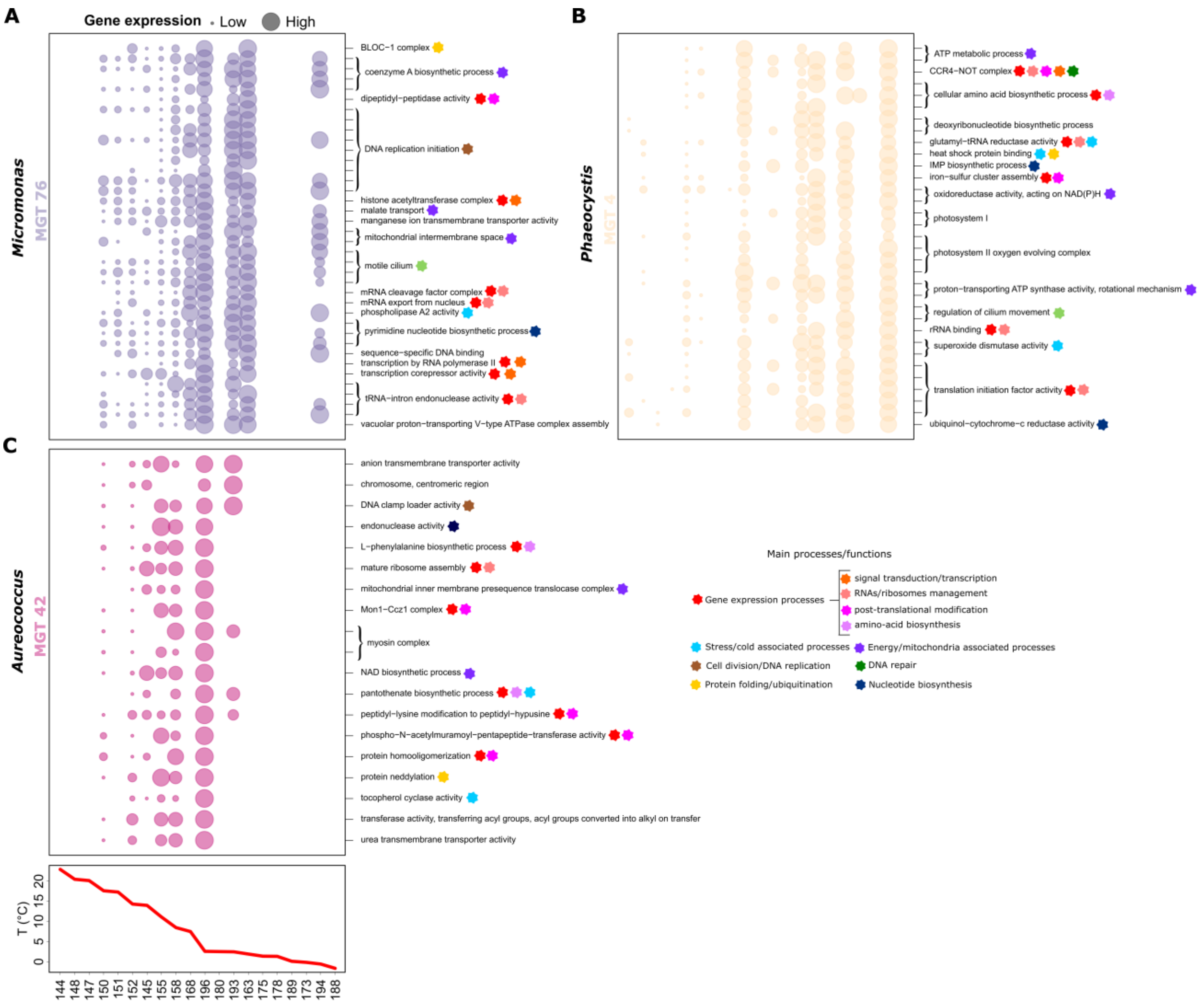

**Figure S16. Expression profiles and Gene Ontology annotation of enriched unigenes negatively correlated with temperature for three algal metagenomics-based transcriptomes (MGTs).** Expression profiles of unigenes whose Gene Ontology terms (right) are enriched within unigenes significantly negatively correlated with temperature are displayed for three algal taxa from three metagenomics-based transcriptomes (MGTs): (**A**) *Micromonas* (MGT 76) (**B**) *Phaeocystis* (MGT 4). For MGT 4, unigenes associated to the 'ribosome' GO term were also enriched but removed from the figure due to their high number (n=24). (**C**) *Aureococcus* (MGT 42). Stations are ordered along the temperature gradient shown at the bottom.

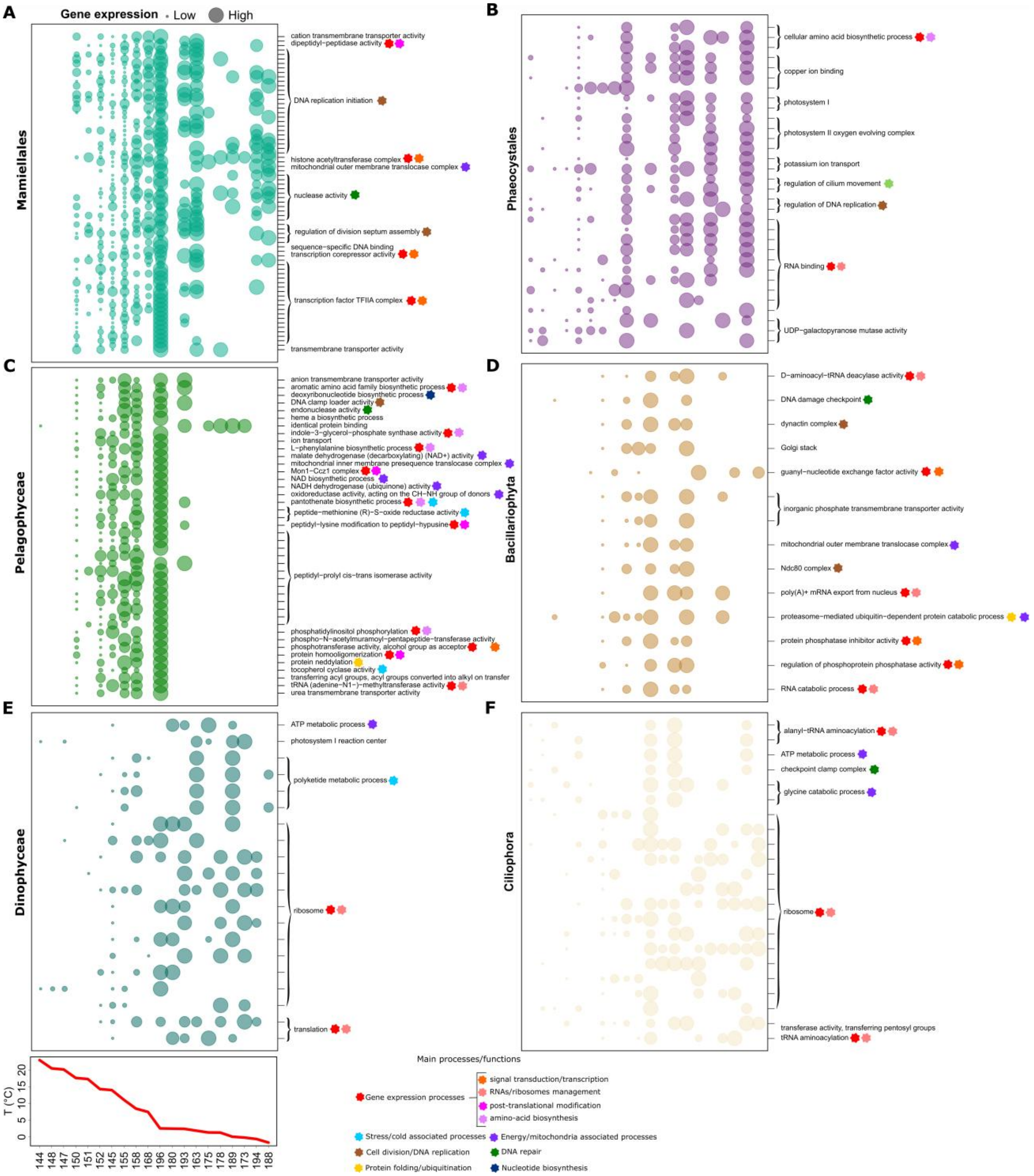

**Figure S17. Expression profiles and Gene Ontology (GO) annotation of enriched unigenes negatively correlated with temperature for six algal taxa.** Expression profiles of unigenes whose Gene Ontology terms (right) are enriched within unigenes significantly negatively correlated with temperature are displayed for six algal taxa: (**A**) Mamiellales (unigenes associated to the 'nucleus' GO term were also enriched but removed from the figure due to their high number (n=27) as well as the unknown, (**B**) Phaeocystales (unigenes associated to the 'ribosome' GO term were also enriched but removed from the figure due to their high number (n=40)) (**C**) Pelagophyceae (**D**) Bacillariophyta (**E**) Dinophyceae and (**F**) Ciliophora. Stations are ordered according to the temperature gradient shown at the bottom.

| | Significantly more abundant functions | |
|---|---|---|
| **Location** | **Metagenomes** | **Metatranscriptomes** |
| **AO** | "cellulose binding"<br>"sphingolipid metabolic process"<br>"ceramide metabolic process"<br>"polynucleotide dephosphorylation"<br>"polynucleotide 5' dephosphorylation"<br>"Mon1-Ccz1 complex"<br>"polynucleotide phosphatase activity"<br>"polynucleotide 5'-phosphatase activity"<br>"transcription, RNA-templated"<br>"RNA-directed 5'-3' RNA polymerase activity" | "hydrolase activity, acting on acid carbon-carbon bonds"<br>"cis-trans isomerase activity"<br>"ribonucleoprotein complex biogenesis"<br>"pseudouridine synthase activity"<br>"hydrolase activity, acting on acid carbon-carbon bonds, in ketonic substances"<br>"preribosome, small subunit precursor"<br>"ribosome biogenesis"<br>"anatomical structure homeostasis"<br>"fumarylacetoacetase activity"<br>"snoRNA binding"<br>"telomere organization"<br>"ribonuclease H2 complex"<br>"lipid catabolic process"<br>"telomeric DNA binding"<br>"S-adenosylmethionine-dependent methyltransferase activity"<br>"intrinsic component of organelle membrane"<br>"microtubule anchoring"<br>"methyltransferase complex"<br>"macromolecule methylation"<br>"integral component of organelle membrane"<br>"vacuole"<br>"sphingolipid metabolic process"<br>"urea metabolic process"<br>"microbody"<br>"tRNA methyltransferase complex"<br>"peroxisome"<br>"tRNA (m1A) methyltransferase complex"<br>"protein transmembrane transport"<br>"chromosome, telomeric region"<br>"ceramide metabolic process"<br>"vacuolar part"<br>"nucleic acid phosphodiester bond hydrolysis"<br>"mitochondrial RNA metabolic process"<br>"peptidyl-amino acid modification"<br>"RNA processing"<br>"RNA modification"<br>"rRNA metabolic process"<br>"peptidyl-proline modification"<br>"protein peptidyl-prolyl isomerization"<br>"pseudouridine synthesis"<br>"ncRNA processing"<br>"serine-type exopeptidase activity"<br>"nuclease activity"<br>"nucleolus"<br>"vacuolar membrane"<br>"urea catabolic process"<br>"RNA methylation"<br>"serine-type carboxypeptidase activity"<br>"endonuclease activity"<br>"tRNA processing"<br>"intracellular protein transmembrane transport"<br>"peptidyl-prolyl cis-trans isomerase activity"<br>"tRNA modification"<br>"tRNA dihydrouridine synthesis"<br>"rRNA processing"<br>"telomere maintenance"<br>"nuclear chromosome, telomeric region"<br>"vacuolar proton-transporting V-type ATPase complex"<br>"RNA methyltransferase activity"<br>"tRNA methylation"<br>"mitochondrial inner membrane presequence translocase complex"<br>"transcription by RNA polymerase I"<br>"tRNA dihydrouridine synthase activity"<br>"tRNA methyltransferase activity" |

| | | |
|---|---|---|
| | | "tRNA (adenine) methyltransferase activity"<br>"tRNA (adenine-N1-)-methyltransferase activity" |
| **NAO** | "actin filament-based process"<br>"pigment metabolic process"<br>"ferrochelatase activity"<br>"glycosaminoglycan binding"<br>"transferase activity, transferring aldehyde or ketonic groups"<br>"intramolecular oxidoreductase activity"<br>"ligase activity, forming carbon-carbon bonds"<br>"ubiquitin-like protein binding"<br>"peptide binding"<br>"pigment biosynthetic process"<br>"ligase activity, forming nitrogen-metal bonds"<br>"unfolded protein binding"<br>"regulation of anatomical structure size"<br>"3-deoxy-7-phosphoheptulonate synthase activity"<br>"argininosuccinate synthase activity"<br>"hydroxymethylbilane synthase activity"<br>"signal sequence binding"<br>"1-deoxy-D-xylulose-5-phosphate synthase activity"<br>"CoA carboxylase activity"<br>"phosphotransferase activity, carboxyl group as acceptor"<br>"aldehyde-lyase activity"<br>"racemase and epimerase activity, acting on carbohydrates and derivatives"<br>"intramolecular oxidoreductase activity, transposing S-S bonds"<br>"protein binding, bridging"<br>"regulation of actin filament-based process"<br>"ubiquitin binding"<br>"ligase activity, forming nitrogen-metal bonds, forming coordination complexes"<br>"cofactor transport"<br>"maintenance of location"<br>"supramolecular fiber organization"<br>"2-isopropylmalate synthase activity"<br>"GMP synthase (glutamine-hydrolyzing) activity"<br>"acetyl-CoA carboxylase activity"<br>"chorismate synthase activity"<br>"fatty acid synthase activity"<br>"fructose-bisphosphate aldolase activity"<br>"glucose-6-phosphate isomerase activity"<br>"phosphoribosylaminoimidazolecarboxamide formyltransferase activity"<br>"ribose-5-phosphate isomerase activity"<br>"mannosamine metabolic process"<br>"cytoskeleton organization"<br>"cellulase activity"<br>"positive regulation of metabolic process"<br>"magnesium chelatase activity"<br>"regulation of organelle organization"<br>"positive regulation of cellular component biogenesis"<br>"ER retention sequence binding"<br>"N-acylglucosamine-6-phosphate 2-epimerase activity"<br>"positive regulation of cellular process"<br>"coenzyme transport"<br>"meiotic cell cycle"<br>"ubiquitin-like protein ligase activity"<br>"uridylyltransferase activity"<br>"regulation of supramolecular fiber organization"<br>"protein disulfide isomerase activity"<br>"IMP cyclohydrolase activity"<br>"3-oxoacyl-[acyl-carrier-protein] synthase activity"<br>"clathrin-coated pit"<br>"N-acetylmannosamine metabolic process"<br>"oxidoreductase activity, acting on the aldehyde or oxo group of donors"<br>"snRNA binding" | "transposition"<br>"transposase activity"<br>"chlorophyll binding"<br>"photosynthetic electron transport chain"<br>"ion channel complex"<br>"photosynthetic electron transport in photosystem II"<br>"electron transporter, transferring electrons within the cyclic electron transport pathway of photosynthesis activity"<br>"protein splicing"<br>"transposition, DNA-mediated"<br>"single strand break repair"<br>"deoxyribonucleotide catabolic process" |

| | |
|---|---|
| | "photosynthesis, light reaction"<br>"light-harvesting complex"<br>"cytokinetic process"<br>"regulation of cytoskeleton organization"<br>"alanine-tRNA ligase activity"<br>"fermentation"<br>"phosphate ion transport"<br>"positive regulation of macromolecule metabolic process"<br>"oxidoreductase activity, acting on the aldehyde or oxo group of donors, NAD or NADP as acceptor"<br>"oxidoreductase activity, acting on the aldehyde or oxo group of donors, disulfide as acceptor"<br>"oxidoreductase activity, acting on diphenols and related substances as donors, oxygen as acceptor"<br>"U6 snRNA binding"<br>"actin cytoskeleton organization"<br>"regulation of protein polymerization"<br>"regulation of cellular component size"<br>"DNA helicase complex"<br>"glycopeptide alpha-N-acetylgalactosaminidase activity"<br>"maintenance of protein location"<br>"negative regulation of cellular component organization"<br>"positive regulation of cellular component organization"<br>"positive regulation of nitrogen compound metabolic process"<br>"maintenance of location in cell"<br>"oxidoreductase activity, acting on CH or CH2 groups, with an iron-sulfur protein as acceptor"<br>"oxidoreductase activity, acting on diphenols and related substances as donors, with copper protein as acceptor"<br>"regulation of microtubule-based movement"<br>"regulation of cilium movement"<br>"glycerol-3-phosphate dehydrogenase [NAD+] activity"<br>"methylenetetrahydrofolate reductase (NAD(P)H) activity"<br>"isoprenoid metabolic process"<br>"O-methyltransferase activity"<br>"S-methyltransferase activity"<br>"regulation of Toll signaling pathway"<br>"plastoquinol--plastocyanin reductase activity"<br>"alternative oxidase activity"<br>"regulation of actin cytoskeleton organization"<br>"5-methyltetrahydrofolate-dependent methyltransferase activity"<br>"4-hydroxy-3-methylbut-2-en-1-yl diphosphate synthase activity"<br>"terpenoid metabolic process"<br>"actin filament organization"<br>"acetyl-CoA carboxylase complex"<br>"photosynthetic electron transport chain"<br>"positive regulation of organelle organization"<br>"negative regulation of organelle organization"<br>"nitrogenous compound fermentation"<br>"positive regulation of cellular metabolic process"<br>"glucose 6-phosphate metabolic process"<br>"alditol phosphate metabolic process"<br>"positive regulation of supramolecular fiber organization"<br>"glycerol-3-phosphate metabolic process"<br>"glutamyl-tRNA reductase activity"<br>"peptidoglycan turnover"<br>"photosynthetic electron transport in photosystem II"<br>"proteasome accessory complex"<br>"chorismate metabolic process"<br>"regulation of actin filament organization"<br>"inorganic phosphate transmembrane transporter |

| | |
|---|---|
| | activity"<br>"small GTPase mediated signal transduction"<br>"positive regulation of protein complex assembly"<br>"electron transporter, transferring electrons within the cyclic electron transport pathway of photosynthesis activity"<br>"positive regulation of cytoskeleton organization"<br>"cell septum assembly"<br>"porphyrin-containing compound metabolic process"<br>"phosphoglycerate kinase activity"<br>"fatty acid metabolic process"<br>"isoprenoid biosynthetic process"<br>"clathrin coat"<br>"positive regulation of protein polymerization"<br>"maintenance of protein location in cell"<br>"phycobilisome"<br>"clathrin-coated vesicle"<br>"heme metabolic process"<br>"Arp2/3 protein complex"<br>"leucine metabolic process"<br>"terpenoid biosynthetic process"<br>"COPI-coated vesicle"<br>"regulation of actin filament length"<br>"heme a metabolic process"<br>"glycerol-3-phosphate catabolic process"<br>"maintenance of protein localization in organelle"<br>“organic anion transmembrane transporter activity"<br>"metalloendopeptidase activity"<br>"methionine metabolic process"<br>"dipeptidyl-peptidase activity"<br>"maintenance of protein localization in endoplasmic reticulum"<br>"L-lysine metabolic process"<br>"metal ion transmembrane transporter activity"<br>"protein polyubiquitination"<br>"protein retention in ER lumen"<br>"regulation of actin polymerization or depolymerization"<br>"trans-Golgi network transport vesicle"<br>"clathrin coat of coated pit"<br>"biotin transport"<br>"exonuclease activity, active with either ribo- or deoxyribonucleic acids and producing 5'-phosphomonoesters"<br>"tail-anchored membrane protein insertion into ER membrane"<br>"fatty acid biosynthetic process"<br>"sulfur amino acid biosynthetic process"<br>"porphyrin-containing compound biosynthetic process"<br>"aspartate family amino acid catabolic process"<br>"exodeoxyribonuclease activity"<br>"monocarboxylic acid transmembrane transporter activity"<br>"clathrin-coated vesicle membrane"<br>"ubiquitin protein ligase activity"<br>"lysine catabolic process"<br>"leucine biosynthetic process"<br>"anaerobic amino acid catabolic process"<br>"exodeoxyribonuclease activity, producing 5'-phosphomonoesters"<br>"COPI-coated vesicle membrane"<br>"heme biosynthetic process"<br>"chlorophyll biosynthetic process"<br>"heme a biosynthetic process"<br>"methionine biosynthetic process"<br>"L-lysine catabolic process"<br>"coenzyme A metabolic process"<br>"pentose-phosphate shunt"<br>"trans-Golgi network transport vesicle membrane"<br>"positive regulation of actin filament polymerization"<br>"pentose-phosphate shunt, non-oxidative branch" |

| | |
|---|---|
| | "clathrin vesicle coat"<br>"GMP metabolic process"<br>"COPI vesicle coat"<br>"queuosine salvage"<br>"methionine synthase activity"<br>"actin nucleation"<br>"L-lysine catabolic process to acetate"<br>"Arp2/3 complex-mediated actin nucleation"<br>"guanosine-containing compound biosynthetic process"<br>"alanyl-tRNA aminoacylation"<br>"clathrin coat of trans-Golgi network vesicle"<br>"biotin transmembrane transporter activity"<br>"GMP biosynthetic process" |

**Table S1. List GO terms found to be significantly more abundant in metagenomes and metatranscriptomes of both the North Atlantic Ocean (NAO) and Arctic Ocean (AO).**

| PFAM | Molecular function and structure | Relevance to cold/hot stress acclimation/response | Refs |
|---|---|---|---|
| PF14169: Cold−inducible protein YdjO | Molecular: Unknown<br>Structure (example of YdjO) [S5]:<br>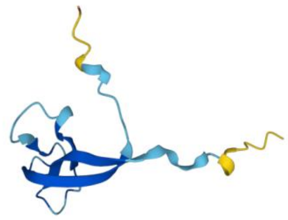 | Strongly cold-shock-induced in a transcriptome analysis of *B. subtilis* | [S6] |
| PF05562: WCOR413 | Molecular: putative transport of substances or ions, mediation of protein/protein interaction<br>Structure: Highly hydrophobic protein with six transmembrane regions | A low-temperature-regulated protein from wheat resembling (25% to 56% identity) a stress-responsive gene from the resurrection plant Xerophyta viscosa Baker (XVSAP1).<br>Hypothesis: "*It is possible that XVSAP1 may be involved in the transport of substances or ions across the plasma membrane as a region stretching from amino-terminal glycine residue of a growing polypeptide. Proteins so modified have diverse functions and the myristate appears to be critical for mediating protein/ protein and/or protein/membrane interactions"* [S7] | [S7,S8] |
| PF00313: CSD | Molecular: Part of this domain is highly similar to the RNP-1 RNA-binding motif [S9].<br>Structure: Protein domain of about 70 amino acids which has been found in prokaryotic and eukaryotic DNA-binding proteins [S9]. | "*may have a function in activating transcription or unwinding or masking RNA molecules.*" [S10] | [S10] |
| PF02649: GCHY−1 | Molecular: tetrahydrofolate biosynthesis. type I GTP cyclohydrolase<br>Structure: highly conserved glutamate residue at position 216 in folE2: likely to be the substrate ligand. The metal ligand is likely to be the cysteine at position 147. $Zn^{2+}$ dependent [S11]. | Cold-upregulated in a thermophilic alga: "*Glycine decarboxylase, which feeds methylene-tetrahydrofolate into the folate cycle was 1.7-fold upregulated at 28°C. In contrast, transcript frequency for formate-tetrahydrofolate ligase, which would drain the folate cycle, was reduced to 0.3-fold at 28°C*" [S12]<br>"*Both* in vitro *and* in vivo *experiments indicate that OtNOS, unlike mammalian NOS, efficiently uses tetrahydrofolate as a cofactor in Arabidopsis plants. The modulation of NO production to alleviate abiotic stress disturbances in higher plants highlights the potential of genetic manipulation to influence NO metabolism as a tool to improve plant fitness under adverse growth conditions."* [S13] | [S11–S13] |
| PF14249: Tocopherol_cycl | Molecular:tocopherol cyclase activity. Secondary metabolism<br>Structure: 3 β-feuillet and 1 α-helix, strong confidence except the tip of the β and α [S5] (N and C-terminal domains) | Abstract: *"As expected, the synthesis of tocopherols (vitamin E) from 2,3-dimethyl-5-phytyl-1,4-benzoquinol by the recombinant protein demonstrated that the EgSXD1 gene product is a TC. Northern hybridizations with RNA isolated from cell suspension cultures clearly showed a fast EgSXD1 downregulation by cold."* [S14] | [S14] |
| PF01964: Thiamine biosynthesis | Molecular: Thiamine biosynthesis, *"Escherichia coli K-12 synthesizes thiamine pyrophosphate (vitamin* | **"***The expression of Arabidopsis genes involved in the thiamine diphosphate biosynthesis pathway, including that of THI1, THIC, TH1 and TPK, was analyzed for 48 h in seedlings* | [S15,S16] |

| | | | |
|---|---|---|---|
| | *B1) de novo. Two precursors [4-methyl-5-(beta-hydroxyethyl)thiazole monophosphate and 4-amino-5-hydroxymethyl-2-methylpyrimidine pyrophosphate] are coupled to form thiamine monophosphate, which is then phosphorylated to make thiamine pyrophosphate."* [S15]<br>Structure: A β-feuillet surrounded by α-helixes and α-β-α on the side (C-terminal) [S16] | *subjected to NaCl or sorbitol treatment. These genes were found to be predominantly up-regulated in the early phase (2-6 h) of the stress response."* [S15] | |
| PF02358: Trehalose_PPase | Molecular: Trehalose phosphatase<br>Structure (example of Trehalose phosphatase) [S5]:<br>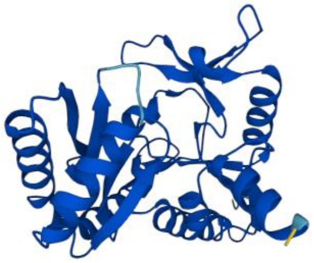 | Abstract: *"A mutant Escherichia coli strain unable to produce trehalose died much faster than the wild type at 4°C. Transformation of the mutant with the otsA/otsB genes, responsible for trehalose synthesis, restored trehalose content and cell viability at 4°C. After temperature downshift from 37°C to 16°C ("cold shock"), trehalose levels in wild-type cells increased up to 8-fold."* [S17] | [S17] |
| PF01979: Amidohydro_1 | Molecular: Allantoinase, Catalyzes the conversion of allantoin (5-ureidohydantoin) to allantoic acid by hydrolytic cleavage of the five-member hydantoin ring [S18].<br>Structure (example of allantoinase) [S5]:<br>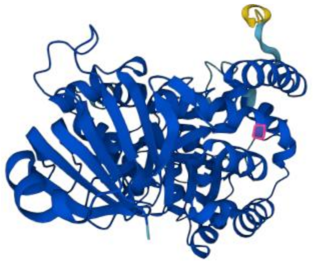 | *"Allantoin is a metabolic intermediate of purine catabolism that often accumulates in stressed plants."* [S19] | [S19,S20] |
| PF10551: MULE | Molecular: Mutator-like element (MULE) transposases. Zn dependent [S21]<br>Structure: This domain was identified by Babu and colleagues [S21]. "*Typical structural features of MULEs include long TIRs (>100 bp) and 8–10 bp flanking* | [S22] | [S21,S22] |

| | | | |
|---|---|---|---|
| | *TSDs*(*63*)” | | |
| PF01119: DNA_mis_repair | Function : DNA repair [S23].<br>Structure: “*This family represents the C-terminal domain of the mutL/hexB/PMS1 family. This domain has a ribosomal S5 domain 2-like fold.*” [S23] | Title: DNA Damage Inducible Protein 1 is Involved in Cold Adaption of Harvested Cucumber Fruit [S24] | [S23,S24] |
| PF00626: Gelsolin | Function :<br>Gelsolin is a cytoplasmic, calcium-regulated, actin-modulating protein that binds to the barbed ends of actin filaments, preventing monomer exchange (end-blocking or capping). It can promote nucleation (the assembly of monomers into filaments), as well as sever existing filaments [S25].<br>Structure (example of Gelsolin) [S5]: uncertain<br>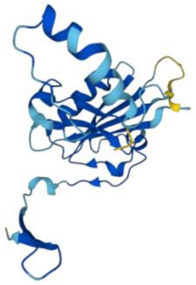 | *“Cold-acclimation had a negligible effect on steady state protein expression in the heart, with just 7 differentially abundant (DA) proteins (0.5% of the quantified cardiac proteome). Furthermore, only gelsolin isoform X1 (log2FC = -0.289) was not influenced by a confounding effect of available development stage”* | [S25] |
| PF00241: Cofilin_ADF | Function:<br>“*They bind both actin-monomers and filaments and promote rapid filament turnover in cells by depolymerising/fragmenting actin filaments. ADF/cofilins bind ADP-actin with higher affinity than ATP-actin and inhibit the spontaneous nucleotide exchange on actin monomers*” [S26]<br>Structure: “*In molecular biology, ADF-H domain (actin-depolymerising factor homology domain) is an approximately 150 amino acid motif that is present in three phylogenetically distinct classes of eukaryotic actin-binding proteins.*” [S27] | *“Based on the above information and on the identification of a wheat cold-regulated ADF homolog, we designed the present experiments to determine the function and regulation of TaADF.”* [S28] | [S26–S28] |
| PF02209: | Molecular: | *“Compared with Table I, only eight spots, which represents seven* | [S29– |

| | | | |
|---|---|---|---|
| VHP | Bundles, nucleates, caps, and severs actin in a Ca2+-dependent manner [S29].<br>Structure: Villin headpiece domain.<br>"*The core of the GH domain consists of a five-stranded b-sheet sandwiched between one long and one short helix*" [S30] | *proteins (38.9%) including a villin (Os03g24220), an actin 3 (Os03g61970), an actin 7 (Os05g01600), an aminotransferase (Os08g41990), an ATP-dependent Clp protease (Os04g32560), a putative subtilisin (Os02g53860), and a sucrose synthase (Os03g28330), showed similar up- or down-regulated patterns."* [S31] | S31] |
| PF03259:<br>Robl_LC7 | Molecular: associated to flagellar outer arm of dynein.<br>Structure:<br>See Koonin et al.[S32] | This family includes proteins that may modulate specific dynein functions, but it may also play a structural or regulatory role. Dynein were shown to be regulated in cold conditions and act as a *"thermally tunable motor"* [S33] | [S32,S33] |
| PF00249:<br>Myb_DNA−binding | Molecular: Myb-like DNA-binding domain (transcription factor)<br>Structure [S5]:<br>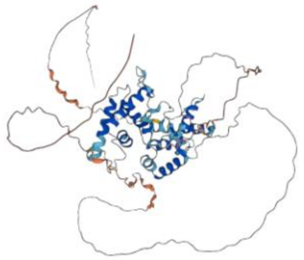 | Putative role in oxidative stress response in *Arabidopsis Thaliana*: regulation of ROS in a circadian manner (probable regulation by CCA1) [S34] | [S34] |
| PF11999:<br>Ice_binding | Molecular: Ice binding, antifreeze glycoproteins, they bind to crystals of ice<br>Structure (example of Ice binding protein 1) [S5]:<br>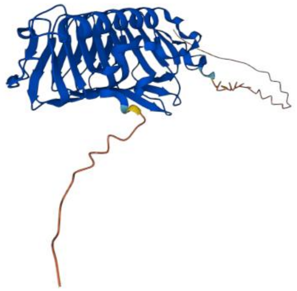 | "*Sub-zero temperatures put plants at risk of damage associated with the formation of ice crystals in the apoplast. Some freeze-tolerant plants mitigate this risk by expressing ice-binding proteins (IBPs), that adsorb to ice crystals and modify their growth.*" [S35] | [S35] |
| PF03104:<br>DNA_pol_B_exo1 | Molecular: 3' to 5' exonuclease activity<br>Structure:<br>ribonuclease H type fold | *"Previous studies [15] showed changes in RNase activity in wheat leaves and roots following exposure to low temperatures (2 8C) for long periods (60 days)."* [S36] | [S36] |
| PF01346:<br>FKBP_N | Molecular: peptidyl-prolyl isomerase (foldase/chaperone), post translation modification<br>Structure [S37]: Domain amino terminal to FKBP-type | "*Some peptidyl prolyl isomerase have been shown to be important in protein folding under cold stress condition by increasing the rate of cis-trans isomerization*" [S38] | [S38] |

| | | | |
|---|---|---|---|
| PF05035:<br>DGOK | Molecular: 2-dehydro-3-deoxygalactonokinase activity<br>Structure (example of deoxygalactonokinase) [S5]:<br>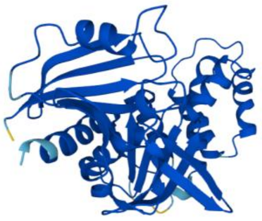 | *"Among these top pathways, the carbohydrate metabolism pathway, galactose metabolism, fructose and mannose metabolism were associated with many up/downregulated genes."* [S39] | [S39] |
| PF05351:<br>GMP_PDE_delta | Molecular:<br>*"PDE delta subunit is thought to be a specific soluble transport factor for certain prenylated proteins and Arl2-GTP a regulator of PDE-mediated transport."* [S40]<br>Structure (example of GMP_PDE_delta containing protein) [S5]:<br>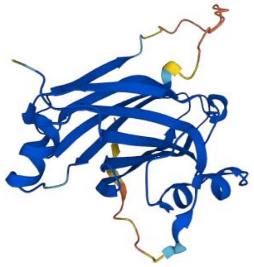 | *"The enzyme is stable only under specific conditions, and the activation property of the enzyme is lost relatively easy. Irreversible modifications occur at temperatures below 0° and above 30°C, and at pH below 6.0. Several other conditions such as high ion concentrations, temperatures just above 0°C and pH above 8.0 lead to reversible modifications of enzyme activity."* [S41] | [S41] |
| PF01180:<br>DHO_dh | Molecular:<br>cytosolic dihydroorotate dehydrogenase<br>Structure (example of dihydroorotate dehydrogenase) [S5]<br>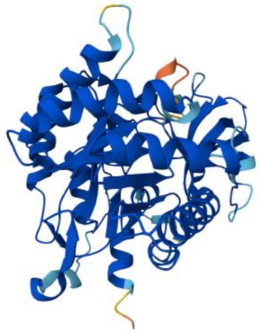 | *"In the present paper, we identified and cloned OsDHODH1 encoding a putative cytosolic dihydroorotate dehydrogenase (DHODH) in rice. Expression analysis indicated that OsDHODH1 is upregulated by salt, drought and exogenous abscisic acid (ABA), but not by cold"* [S42]<br>*"Even though there is no data on the effect of temperature on DhoDH, we can expect that its thermal sensitivity will affect more mitochondrial functions than only the reduction of the Q pool."* [S43] | [S42,S43] |
| PF05971:<br>Methyltransf_10 | Molecular:<br>RNA methyltransferase<br>Structure (example of Methyltransferase) [S5]: | RNA methylation is known to be important in stress response especially cold acclimation (stability of RNA) [S44] | [S44] |

| | |
|---|---|
| | 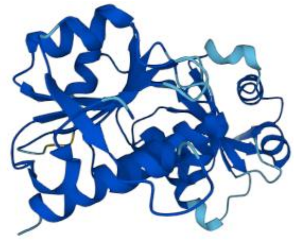 |

**Table S2. Summary table of function and structure of a set of PFAMs known to be involved in cold or hot stress acclimation.**

| Metabolic function | Sub function | PFAMs |
|---|---|---|
| Photosynthesis (P) | Electron transport chain (Main) | PF00127, PF02276, PF01716, PF00223, PF02605, PF07465, PF05479, PF00737, PF02532, PF02533, PF05151, PF02468, PF04725, PF01405, PF06514, PF07123, PF06596, PF06298, PF05969, PF13326, PF00796, PF02427, PF02507, PF03244, PF01701, PF01241, PF10657, PF11947 |
| | Light Harvest (LH) | PF00504, PF00556 |
| | Cyclic electron transport (CET) | PF00124, PF00421, PF03967, PF11623 |

**Table S3. List of PFAMs used in the analysis of photosynthesis-related functions.**
